# Aligning biological sequences by exploiting residue conservation and coevolution

**DOI:** 10.1101/2020.05.18.101295

**Authors:** Anna Paola Muntoni, Andrea Pagnani, Martin Weigt, Francesco Zamponi

## Abstract

Aligning biological sequences belongs to the most important problems in computational sequence analysis; it allows for detecting evolutionary relationships between sequences and for predicting biomolecular structure and function. Typically this is addressed through profile models, which capture position-specificities like conservation in sequences, but assume an independent evolution of different positions. RNA sequences are an exception where the coevolution of paired bases in the secondary structure is taken into account. Over the last years, it has been well established that coevolution is essential also in proteins for maintaining three-dimensional structure and function; modeling approaches based on inverse statistical physics can catch the coevolution signal and are now widely used in predicting protein structure, protein-protein interactions, and mutational landscapes. Here, we present DCAlign, an efficient approach based on an approximate message-passing strategy, which is able to overcome the limitations of profile models, to include general second-order interactions among positions and to be therefore universally applicable to protein- and RNA-sequence alignment. The potential of our algorithm is carefully explored using well-controlled simulated data, as well as real protein and RNA sequences.

## I. INTRODUCTION

In the course of evolution, proteins undergo substantial changes in their amino-acid sequences, while keeping their three-dimensional fold structure and their biological function remarkably conserved. In computational biology, this structural and functional conservation is extensively used: when we can establish that two proteins are homologous, i.e. they share some common evolutionary ancestor, properties known for one protein can be used to computationally annotate its homolog. Even at the finer amino-acid scale, two positions detected to be homologous are expected to have the same positioning inside the protein structure, and share common functionality (e.g. active sites or binding interfaces).

Detecting homology is, however, not an easy task. First, homologous proteins may share only 20% or even less of their amino acids, the others being substituted in evolution, making the detection of similarity rather involved. Even worse, proteins may change their length, amino acids may be inserted into a sequence, or deleted from it. Just looking to a single sequence, we have no information on which positions might be insertions or deletions, and which positions might be inherited from ancestral proteins, possibly undergoing amino-acid substitutions.

To solve this problem, *sequence alignments* have to be constructed [1]. The objective of sequence alignments is to identify homologous positions, also called matches, along with insertions and deletions, such that the aligned sequences become as similar as possible. In this context, three frequently used, but distinct alignment problems can be identified:

- *Pairwise alignments* compare two sequences. Under some simplifying assumptions, cf. below, this problem can be solved efficiently using dynamic programming (i.e. an iterative method similar to transfer matrices or message passing in statistical physics) [2, 3]. Detecting homology by pairwise alignment is limited to rather close homologs.
- More distant homology can be detected using *multiple-sequence alignments* (MSA) of more than two proteins [4], which minimize the global sequence similarities by constructing a rectangular matrix formed by amino acids and gaps, representing both insertions and deletions. The rows of this matrix are the individual proteins, the columns aligned positions. Besides being able to detect more distant homology, MSA allow for identifying conserved positions, i.e. columns which do not (or rarely) change the amino acid. These positions are typically known to be important, either for the functionality (e.g. active sites in proteins) or for the thermodynamic stability of the protein fold. Although dynamic programming methods can be generalized from pairwise to multiple-sequence alignments, their running time becomes exponential in the number of sequences to be aligned. Many heuristic strategies have been proposed following the seminal ideas of [5], cf. [6–9], but the construction of accurate MSA of more than about 10^3^ sequences remains challenging.
- For larger alignments, a simple strategy is widely used. Instead of constructing a globally optimal sequence alignment, *sequences are aligned one by one to a well-constructed seed MSA* [10, 11]. As in the case of MSA, this strategy allows for detecting distant homologs and for exploiting amino-acid conservation. If the seed MSA is reasonably large (*≥*10^2^ sequences) and of high quality, very large and rather accurate alignments of up to 10^6^ sequences can be constructed easily. This strategy is currently the best choice for constructing large families of homologous proteins [12], and is also the subject of our work.

Up to one major exception discussed below, almost all sequence-alignment methods are based on the simplifying assumption of *independent-site evolution*. In this setting, global sequence similarity can be expressed as a sum over similarities of individual columns. In term of sequence statistics, this accounts for assuming that the global probability of observing some full-length sequence can be factorized over site-specific but independent single-site probabilities, also known as *profile models*, cf. [1], which are able to capture amino-acid conservation, but no correlations between positions. A successful variant are *profile Hidden Markov Models* [13], which assume independence of matched positions, but take into account that gap stretches are more likely than many individual gaps to reflect the tendency of homologous proteins to accumulate in the course of evolution large-scale modular gene rearrangements.

A major advantage of profile models is their computational efficiency, as they allow for determining optimal alignments in polynomial time using dynamic programming, cf. [1]. They also allow to take benefit from conserved positions, which serve intuitively as anchoring point in aligning new sequences to the seed MSA. Variable positions contribute much less to profile-based sequence alignment.

An important exception are so-called *covariance models* for functional RNA [14]. RNA sequences are characterized by low sequence conservation, making alignment via profile models unreliable, but highly conserved secondary structures, due to base pairing inside the single-stranded RNA molecules, and the formation of local helices, the so-called stems. Base pairing does not pose constraints on the individual bases, but on the correct pairing in Watson-Crick pairs A:U and G:C, or wobble pairs G:U, which consequently have to be described by a non-factorisable pair distribution. In the case of RNA, the planar structure of the graph formed by the RNA chains and the base-pairings still enable the application of exact but computationally efficient dynamic programming [14–17].

In the last decades, amino-acid covariation, or coevolution, has been established as a statistically important feature of protein evolution [18]. Modeling MSA statistics via methods related to the so-called Direct-Coupling Analysis (DCA) [19–22], inspired by inverse statistical physics [23–26], has found widespread applications in protein-structure prediction from sequence [27–30], detection of protein-protein interactions [19, 31–37], of mutational effects in proteins [38–42] or even in data-driven sequence optimization and design [39, 43, 44]. DCA is based on the assumption of a generalized Potts model with disordered pairwise couplings between sites.

The resulting situation is somewhat paradoxical: pair-wise Potts models are inferred from MSA, but MSA are constructed using an independent-site assumption. Important structural and functional information is contained in the covariation of amino-acids, but it is neglected in the alignment procedures used for proteins.

Our work aims at overcoming this paradox, by including the information contained in amino-acid (or nucleotide) covariation in aligning sequences to a seed MSA. This idea shows important similarity to that of covariance models and RNA alignment, but the lack of planarity of the underlying fully connected DCA models makes an application of dynamic programming impossible. We cope with this problem by proposing an approximate message passing strategy based on Belief Propagation [45], further simplified in a high-connectivity mean-field limit for long-range couplings [46].

The plan of the paper is the following: we first formalize the problem and its statistical-physics description. The latter allows us to derive DCAlign, a combined belief-propagation / mean-field algorithm for aligning a sequence to a Potts model constructed by DCA from the seed MSA. The efficiency of our algorithm is first tested in the case of artificial data, which allow us to evaluate the influence of conservation (i.e. single-site statistics) and covariation (i.e. two-site couplings) in the alignment procedure. Extensive tests and positive results are given for a number of real protein and RNA families. Technical details of the derivation of the algorithm are provided in the Supplementary Information (SI).

## II. SETUP OF THE PROBLEM

### A. Alignment

The method we are going to describe can be applied to align different types of biological sequences (viz. DNA, RNA, proteins). We discuss here the protein case, but the extension to the other cases is straightforward and it will be considered below. Let us consider an amino-acid sequence ***A*** = (*A*_1_, …, *A*_*N*_) containing a protein domain ***S*** = (*S*_1_, …, *S*_*L*_) of a known family, which we want to identify. Note that ***S*** may contain amino acids and gaps, while the original sequence ***A*** is composed exclusively by amino acids. We assume that the protein family is well described by a Direct Coupling Analysis (DCA) model, or Potts Hamiltonian, or simply “energy”,

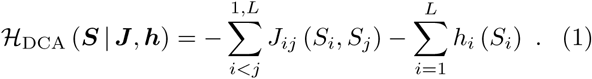

Here, the sequence ***S*** = (*S*_1_, …, *S*_*L*_) is assumed to be aligned to the MSA of length *L* of the protein family, and the set of parameters ***J*** and ***h*** are considered as known, having been learned from some seed alignment [47]. The energy ℋ_DCA_ is then considered as a “score” (lower energy corresponds to higher score) for sequence ***S*** to belong to the protein family. We address the problem of aligning a sequence ***A*** to the model ℋ_DCA_ or, in other words, of detecting the domain in ***A*** that has the best score within the model ℋ_DCA_. In this setting, the solution to our problem is the sub-sequence (cf. below for the precise definition of a sub-sequence including insertions and deletions) that, among all the possible sub-sequences of ***A***, minimizes the energy (or, at finite temperature, is a typical sequence sampled from ℋ_DCA_). The energy thus serves as a cost function for comparing different candidate alignments. Contrarily to profile models, which only take into account conservation of single residues, DCA models also include pairwise interactions related to residue coevolution (and thus in particular at any linear separation along the sequence ***A***), hopefully leading to more accurate alignments in cases where conservation alone is insufficient.

However, the DCA model ℋ_DCA_ does not model insertions, because the parameters ***J*** and ***h*** are inferred from a seed MSA where all columns containing inserts have been removed. A suitable additional cost has thus to be assigned to amino-acid insertions, which are needed in order to find a low-cost alignment. Still, we have to prevent our algorithm from picking up energetically favorable but isolated amino acids out of the (possibly long) input sequence ***A***. For modeling this cost, we will explicitly refer to the insertion statistics in the full seed alignment.

Note that the DCA model contains position-specific gap terms in the ***J*** and ***h***; so the gap statistics of the seed MSA is fully described by the DCA model alone. Nonetheless, the observed statistics deeply depends on how the seed is constructed, and it could a priori be non-representative of the gap statistics of the full alignment. To take into account this degree of variability, we allow for the introduction of an additional energy term associated with the presence of gaps. A more detailed discussion is reported in Sec. II B.

Formally speaking, the alignment problem reduces to finding a sub-sequence ***S*** = (*S*_1_, …, *S*_*L*_) of ***A*** = (*A*_1_, …, *A*_*N*_) such that:

1. the sub-sequence ***S*** forms an ordered list of aminoacids in ***A*** (“match” states) with the possibility of (a) adding gaps, denoted as “–”, between two consecutive positions in ***A*** and (b) skipping some amino acids of ***A***, i.e. interpreting them as insertions;
2. the aligned sequence ***S*** minimizes

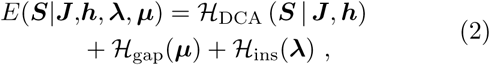

being ℋ_gap_(***µ***) a penalty for adding gaps that depends on the number and position of gaps and on the hyper-parameters ***µ***, and ℋ_ins_(***λ***) being a penalty on insertions, parametrized by the hyper-parameters ***λ***.

We first consider here the case where *N ≳ L*, i.e. we are trying to align a domain in a longer sequence, or *N < L* when we are trying to align a fragment. The case *N ≫ L*, i.e. when we search for a hit of the DCA model in a long sequence, may be computationally hard for the approach proposed here, because the alignment time scales roughly as *L*^2^*N* ^2^, as discussed below. Current state-of-the-art alignment methods like BLAST use heuristics to approximately locate possible hits, and perform accurate alignment search only in these restricted regions, to speed up search. It might be necessary to do this before running DCAlign, but this is not the objective of the current work. In general, it will be better if *N* is not too different from *L*.

An example of a full sequence and its alignment is given in Fig. 1. The sequence on the top is a full sequence of *N* = 19 amino-acids, and the highlighted part is the target domain. The aligned sequence of length *L* = 10, reported on the bottom, consists in one gap in position 2 and 9 matched amino-acids.

**FIG. 1.**
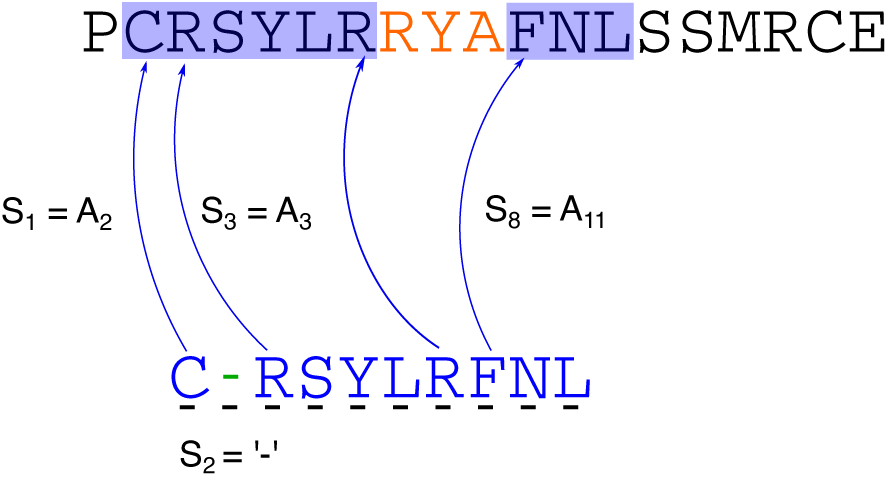
Example of an alignment. On the top, we show a full length sequence ***A***, and on the bottom its alignment ***S***, in which both gap and insertion events occur. The domain to be aligned is highlighted in blue. We show explicitly three matched states *S*_1_ = *A*_2_, *S*_3_ = *A*_3_ and *S*_8_ = *A*_11_ and a gap insertion in *S*_2_ =“–”. In this example we also show a possible way of skipping some amino-acids in the original sequence, that is to assign three insertions, highlighted in red.

### B. Gap and insertion penalties

The hyper-parameters ***µ, λ*** determine the cost of adding a gap or an insertion in the aligned sequence. They must be carefully determined to allow for these events without affecting the quality of the alignment. In other words, we would like to reduce, as much as possible, the number of gaps (when the statistics of gaps of the seed we use is biased) and to parsimoniously add insertions when energetically favorable, avoiding to pick up isolated amino acids in the alignment.

To deal with insertions we use a so-called affine penalty function [1] parametrized by 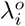, the cost of adding a first insertion between positions *i* − 1 and *i*, and 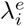, the cost of extending an existing insertion, with 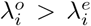. This results in

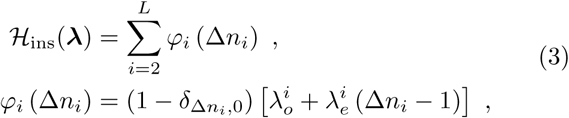

where Δ*n*_*i*_ is the number of insertions between positions *i* − 1 and *i*. This set of parameters can be learned from a seed alignment through a Maximum Likelihood (ML) approach as reported in Sec. IV B. Finally, we introduce two types of gap penalties, denoted by *µ*^int^ and *µ*^ext^, which are associated with an “internal” gap between two matched states and with an “external” gap (at the beginning and at the end of the aligned sequence), respectively. This gives

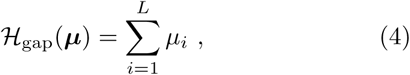

where *µ*_*i*_ = 0 for match states, *µ*_*i*_ = *µ*^int^ for internal gaps and *µ*_*i*_ = *µ*^ext^ for external gaps.

An illustration is given by the aligned sequence in Fig. 1. Insertions are highlighted in red, and the total insertions penalty is then given by 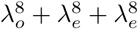. A gap, which increases the total energy by *µ*^int^, is highlighted in green at position 2 of ***S***.

### C. Statistical physics model

We now want to construct a discrete statistical-physics model which defines this alignment. For the positions 1 *≤ i ≤ L*, the model has to encode the position of the gaps and of the match states, with their corresponding symbol in the sequence (*A*_1_, …, *A*_*N*_). We therefore introduce two variables per site 1 *≤ i ≤ L*. The first one is a boolean “spin” *x*_*i*_ *∈* {0, 1}, which tells us if *S*_*i*_ is a gap (*x*_*i*_ = 0) or an amino-acid match (*x*_*i*_ = 1). The second one is a “pointer” *n*_*i*_ *∈* {0, …, *N* + 1}, which gives, for the case of match states *x*_*i*_ = 1 and 1 *≤ n*_*i*_ *≤ N*, the corresponding position in the original sequence (*A*_1_, …, *A*_*N*_); note that this allows for insertions if *n*_*i*+1_ − *n*_*i*_ *>* 1. If matched symbols start to appear only from a position *i >* 1, we then fill the previous positions {*j* : 1 *≤ j < i*} with gaps having pointer *n*_*j*_ = 0. Similarly, if the last matched state appears in *i < L* we fill a stretch of gaps in positions {*j* : *i < j ≤ L*} having *n*_*j*_ = *N* + 1. This encoding allows one to distinguish the “external” gaps at the boundary of the aligned sequence, whose total number we denote as 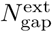, from the “internal” ones, i.e. between two consecutive matched states, whose total number is 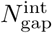. Formally, the number of gaps and insertions are

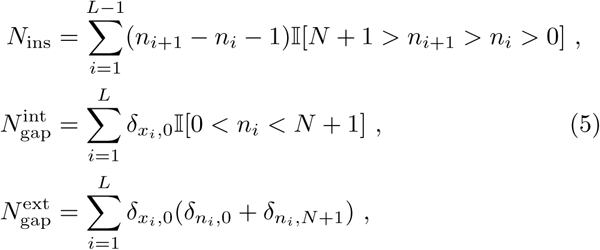

where 𝕀[*ε*] is the indicator function of the event *ε*. Introducing the short-hand notation *A*_0_ = “–” (gap), a model configuration (*x*_1_, …, *x*_*L*_, *n*_1_, …, *n*_*L*_) results in an aligned sequence 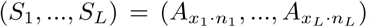 The auxiliary variables (***x, n***) must be additionally assigned such that the positional constraints illustrated in Fig. 1 are satisfied, i.e. the target sub-sequence must be ordered, as we now describe.

First of all, in order to correctly set the pointers in presence of gaps in the first and last positions, it is sufficient to set the state of node *i* = 1 as

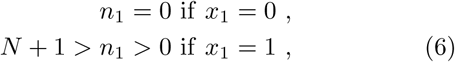

and the state of node *i* = *L* as

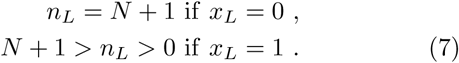

These properties can be formally expressed by the following two single-position constraints

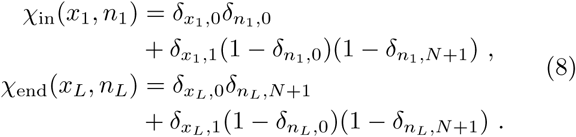

Next, we need to locally impose that, for each position 1 *< i < L*,

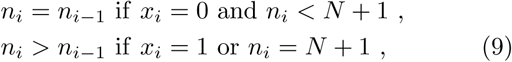

i.e. the pointer *n*_*i*_ remains constant when *x*_*i*_ = 0, and it jumps to any later position in *n*_*i*−1_ + 1, …, *N* if *x*_*i*_ = 1. This jump, besides determining the amino-acid *S*_*i*_ to be placed in position *i*, also allows for identifying inserts according to Eq. (5). A pictorial representation of this constraint is shown in Fig. 2. We can formally encode these constraints in a “short-range” function *χ*_sr_(*x*_*i*−1_, *n*_*i*−1_, *x*_*i*_, *n*_*i*_) that, for each pair of consecutive positions (*i* − 1, *i*), indicates the feasible/unfeasible configurations of the variables (*x*_*i*−1_, *n*_*i*−1_, *x*_*i*_, *n*_*i*_) and the associated cost of insertions, as

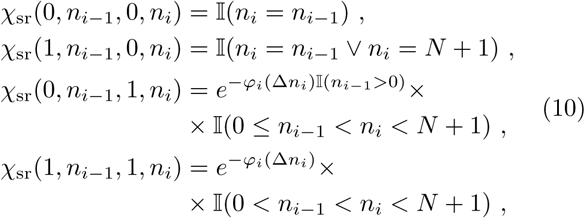

where the function *φ*_*i*_(Δ*n*_*i*_) is the contribution of the *i*−th position to the affine insertion penalty, as given in Eq. (3) with Δ*n*_*i*_ = *n*_*i*_ − *n*_*i*−1_ − 1. Note that combining the constraints in Eq. (8) and Eq. (10), positions {*j >* 1} ({*j < L*}) can either have a gap with *n*_*j*_ = 0 (*n*_*j*_ = *N* + 1) or the first (last) match at any position *n*_*j*_ *>* 0 (*n*_*j*_ *< N* + 1) with no insert penalty.

**FIG. 2.**
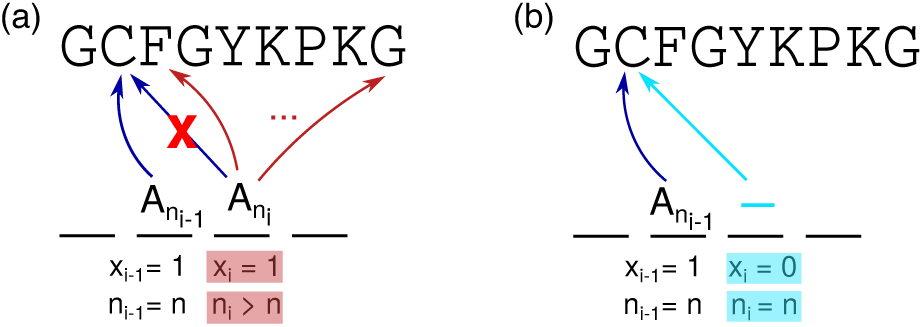
Short range constraints. We plot in (a) a feasible assignment of two consecutive matched states. If in position *i*− 1 we assign *S*_*i*−1_ = *A*_*n*_, we can then align the next position *i* to one of the possible amino-acids *S*_*i*_ *∈* {*A*_*n*+1_, …, *A*_*N*_}. As a consequence (*x*_*i*−1_, *x*_*i*_) = (1, 1), *n*_*i*−1_ = *n* while *n*_*i*_ *∈* {*n* + 1, …, *N*}. In (b) we plot a feasible inclusion of a gap in position *i*. If the previous site *i* − 1 points to *n*_*i*−1_ = *n*, we then assign (*x*_*i*_, *n*_*i*_) = (0, *n*) to keep memory of the aligned sequence. In the next position, *i*+1, we can match an amino-acid in further positions (according to the constraint in (a)) or add another gap with pointer *n*_*i*+1_ = *n*.

Finally, gap penalties can be encoded in a single-variable weight,

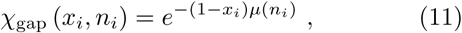

With

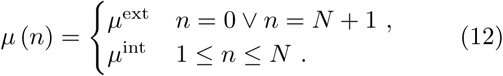

The DCA Hamiltonian can be rewritten in terms of the auxiliary variables as

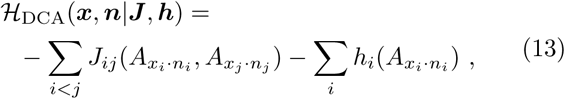

while the global cost function *E* in Eq. (2) is

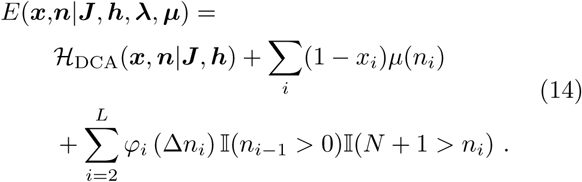

Collecting all these definitions together, we can associate a Boltzmann weight *W* (***x, n***) with each possible alignment (***x, n***) of a given sequence ***A***, which takes into account all energetic contributions for feasible assignments only,

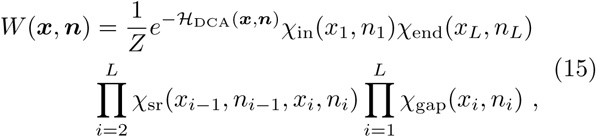

where *Z* is the partition function. Note that the “hard constraints” *χ*_sr_, which can set the weight to zero, live only on the edges of the linear chain 1, …, *L*, while the interactions *J*_*ij*_ are in principle fully connected.

Finally, we can map the original minimization problem in Eq. (2) as the statistical physics problem of finding the best assignment of the variables (***x***^*^, ***n***^*^) that maximizes the Boltzmann distribution:

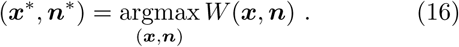

Alternatively, we could obtain an optimal alignment (***x***^*^, ***n***^*^) from an equilibrium sampling of alignments with weight *W* (***x, n***). Unfortunately, both sampling from *W* (***x, n***) and identifying the constrained optimal assignment are hard and intractable problems. Note that the space of possible assignments has dimension scaling as (*N* + 2)^*L*^, which grows extremely quickly with *N* and *L*. For comparison, the DCA problem is defined in a space growing “only” as *q*^*L*^. However, some approximations inspired by statistical physics can be exploited for seeking an approximate solution.

## III. ADVANCED MEAN-FIELD APPROXIMATION

A straightforward approach to make this problem tractable is to use message-passing approximations of the marginal probabilities *P*_*i*_(*x*_*i*_, *n*_*i*_) of Eq. (15), such as Belief Propagation (BP), which are exact for problems defined on graphs without loops. Note that BP is also exact on linear chains, for which it coincides with the transfer matrix method (or dynamic programming / forward-backward algorithm [1]). One can think to BP as treating exactly the linear chain 1, *…, L*, while the longer-range interactions are approximated by message-passing. In the case of vanishing couplings *J*_*ij*_(*A, B*) *=* 0 for all *i, j* such that |*i*, − *j*| *>* 1 and for all *A, B*, i.e. in the case of a model with nearest-neighbor interactions only, this formulation is exact; if all couplings vanish, it is quite similar to a profile Hidden Markov Model, which has the same penalties for opening and extending a sequence of insertions. However, our interactions are instead typically very dense (all couplings are non-zero, hence the associated graph is very loopy), but weak. We can thus consider a further approximation of BP [46], in which the linear chain 1,*…, L* is still treated exactly, while the contribution of more distant sites is approximated via mean field (MF), in a way similar to Thouless-Anderson-Palmer (TAP) equations [48], also known as Approximate Message Passing (AMP) equations. We refer to this approach simply as MF in the following. In the rest of this section we derive the BP and MF equations.

### A. Transfer matrix equations for the linear chain

Suppose first that only nearest-neighbor couplings *J*_*i,i*+1_ (*A, B*) are non-zero. In this case, the problem is exactly solved by the transfer matrix method, which corresponds to a set of recursive equations for the “forward messages” *F*_*i*_(*x*_*i*_, *n*_*i*_) = *F*_*i*→*i*+1_(*x*_*i*_, *n*_*i*_), i.e. the probability distribution of site *i* in absence of the link (*i, i* + 1), and “backward messages” *B*_*i*_(*x*_*i*_, *n*_*i*_) = *B*_*i*→*i*−1_(*x*_*i*_, *n*_*i*_), i.e. the probability distribution of site *i* in absence of the link (*i* − 1, *i*).

We give here the transfer matrix equations for the forward and backward messages. For compactness we define the single-site weight contribution to Eq. (15):

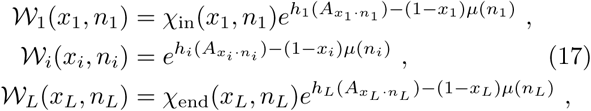

where the second line is for *i* = 2, *…, L* 1. We then have for the forward messages:

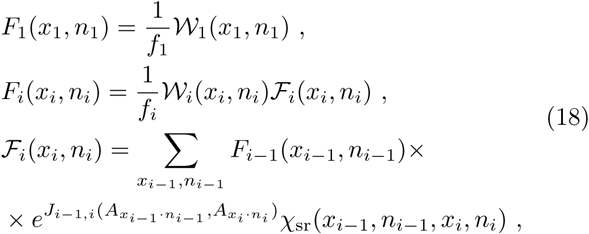

where *F*_*i*_ is defined for *i* = 1, …, *L* − 1 and *F*_*i*_ for *i* = 2,, *L*, and the *f*_*i*_ are normalization constants determined by the requirement that messages are normalized to one. For the backward messages we have

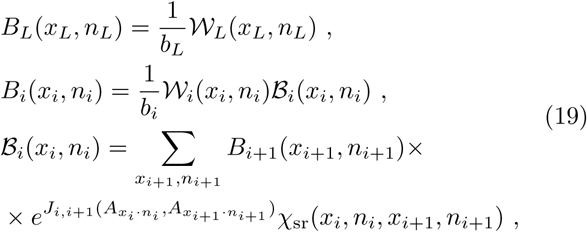

where *B*_*i*_ is defined for *i* = *L, L* − 1, …, 2 and ℬ_*i*_ for *i* = *L* − 1,*…*, 1, and *b*_*i*_ are normalization constants. Finally, the marginal probabilities are given by

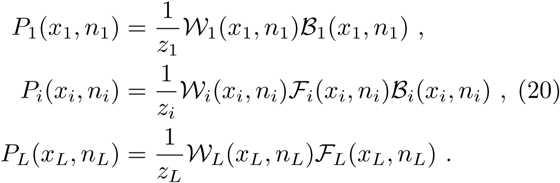

These equations are summarized in compact form in Table I. They can be easily implemented on a computer and solved in a time scaling as *LN*.

**TABLE I.** Schematic summary of the transfer matrix and mean field equations, which are complemented by the recurrence equations for ℱ_*i*_ and ℬ_*i*_ given in Eqs. (18) and (19). For mean field one should replace 𝒲_*i*_ → 𝒞_*i*_.

| $i = 1$ | $i = 2, \dots, L-1$ | $i = L$ |
| --- | --- | --- |
| $P_1 = \frac{1}{z_1} \mathcal{W}_1 \mathcal{B}_1$ | $P_i = \frac{1}{z_i} \mathcal{W}_i \mathcal{F}_i \mathcal{B}_i$ | $P_L = \frac{1}{z_L} \mathcal{W}_L \mathcal{F}_L$ |
| $F_1 = \frac{1}{f_1} \mathcal{W}_1$ | $F_i = \frac{1}{f_i} \mathcal{W}_i \mathcal{F}_i$ | — |
| — | $B_i = \frac{1}{b_i} \mathcal{W}_i \mathcal{B}_i$ | $B_L = \frac{1}{b_L} \mathcal{W}_L$ |

### B. Long range interactions

We now discuss the inclusion of long-range interactions in the transfer matrix scheme. In order to treat correctly the long-range interaction in BP, it is important to note that the same “light-cone” condition expressed by the constraint *χ*_sr_ in Eq. (10) holds between any pair (*i, j*). However, this condition would be violated by the messages of BP due to their approximate character on loopy graphs. In order to enforce it, we can introduce a new constraint

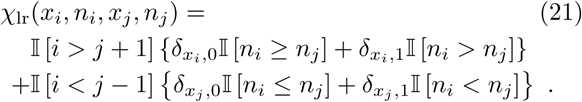

Because this constraint is redundant with respect to Eq. (10), it can be added without changing the weight:

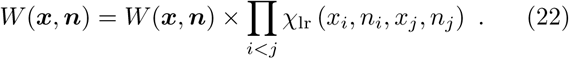

However, adding this constraint ensures that the proper ordering of the pointers is preserved under the BP approximation.

The BP equations can be written straightforwardly (see SI) and provide an approximation to the marginal probabilities *P*_*i*_ in a time scaling as *L*^3^*N* ^2^, which is not very convenient. In the SI we discuss a simplification of the BP equations, under the assumption that pairs of sites with | *i*− *j*| *>* 1 can be treated in a mean field [46]. We find that the resulting mean field equations are identical to the transfer matrix ones, with the only replacement of the local weight 𝒲_*i*_ → 𝒞_*i*_, with

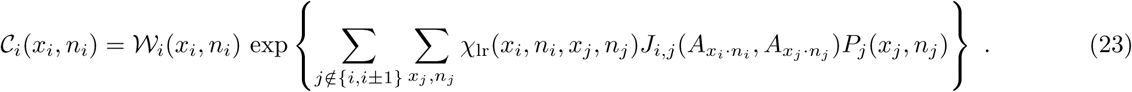

As a result, the mean field equations have the same complexity as the transfer matrix equations (*LN*) with an additional factor *LN* needed to compute each *𝒞*_*i*_, resulting in an overall complexity *L*^2^*N* ^2^.

### C. Assignment

After solution of the MF equations, from the marginal probabilities {*P*_1_(*x*_1_, *n*_1_), …, *P*_*L*_(*x*_*L*_, *n*_*L*_)} we have to find the most probable assignment (***x***^*^, ***n***^*^), as defined in Eq. (16). The simplest way to do so is to assign to each position *i* the most probable state according to its marginal, i.e.

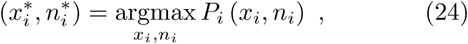

which is possible whenever the obtained assignment satisfies all the hard constraints. However, in some cases, the set of locally optimal positions do not satisfy the short-range constraints due to the approximate nature of the MF solution. We then perform a *maximization* step, in which we select the position *i*^*^ and the local assignment 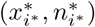 having the largest probability among all the marginals, i.e.

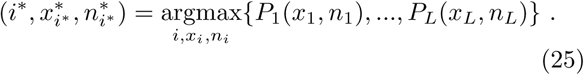

We then set the state of site *i*^*^ in 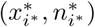, and we proceed with a *filtering* step, in which we set to zero the marginal probabilities of the incompatible states of the first nearest neighbors of *i*^*^. In practice, we multiply the marginals by the short-range constraints computed at 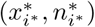, i.e. we consider the new marginals on sites *i*^*^ *±* 1:

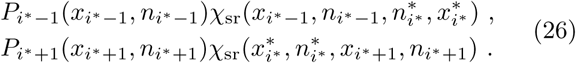

We can now repeat the *maximization* step in order to find the state (*x*^*^, *n*^*^) that maximizes the joint set of probabilities for the positions adjacent to the already aligned part of the sequence (in this case *i*^*^ −1 and *i*^*^ +1, because we only fixed *i*^*^),

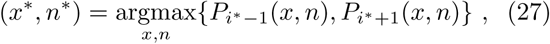

and we fix this state, in the alignment, in the right position (either *i*^*^ + 1 or *i*^*^ 1). Suppose for simplicity that we have just specified the state in position *i*^*^ + 1; we now filter the probability of *i*^*^ +2 and repeat the choice for the next (*x*^*^, *n*^*^) considering the set of (modified) marginals {*P*_*i*\*−1_(*x, n*), *P*_*i*\*+2_(*x, n*)}. The procedure is repeated until all the *L* positions are determined.

Note that this scheme is somehow greedy, because the assignments are decided step by step, and are constrained by the choices made in the previous positions. Still, the assignment is guided by the marginal probabilities obtained from considering the global energy function and all the hard constraints. Moreover, this assignment procedure is as fast as the “max-marginals” scheme because it does not require to re-run the update of the equations in Sec. III, and thanks to the step-by-step filtering of the marginals it ensures an outcome that is always compatible with the constraints.

### D. Discussion

In this section we presented a set of approximate equations and an assignment procedure that, together, allow us to solve the alignment problem in polynomial time. The BP equations have a computational complexity proportional to *L*^3^*N* ^2^, while the MF equations scale as *L*^2^*N* ^2^. In both cases, the equations can be solved at “temperature” equal to one, corresponding to a Boltzmann equilibrium sampling from the weight in Eq. (15), or at zero temperature, corresponding to finding (approximately) the most likely assignment in Eq. (15) (the full set of equations at zero temperature is reported in the SI). Of course, any intermediate temperature could also be considered, but we do not explore other values of temperature in this work.

In all cases, one can compute the free energy associated with the BP or MF solution, which gives a “score” measuring the quality of the alignment. This score could be used, in long sequences with multiple hits, to decide a “best hit”. The expression of the free energy is given in the SI.

The MF equations are derived from the BP equations by assuming that all couplings with |*i* − *j*| *>* 1 are weak enough to be treated in mean field. But we know that in (good) protein models, stronger couplings correspond to physical contacts in the three-dimensional structure, while a background of weaker couplings describe other correlations or even just noise. It could be interesting, therefore, to use a mixed BP/MF method, in which weaker couplings with ∥*J*_*ij*_∥ *< K* are treated in mean field, while stronger couplings with ∥*J*_*ij*_ ∥ *≥ K* are treated with BP, for a given threshold *K*. The case *K* = 0 corresponds to pure BP, while the case *K* → *∞* corresponds to pure MF. One could check whether an optimal value of *K* exists. We leave this for future work.

## IV. LEARNING THE MODEL

We now discuss the learning of the model parameters, namely the couplings and fields of the DCA Hamiltonian, and the hyper-parameters ***µ, λ***.

### A. Potts model

Our alignment method is able to cope with different cost functions, because the implementation of the update equations described in Sec. III B is as general as possible. Introducing a 5-state alphabet, the method is also able to treat RNA alignments.

In this work, we tested several types of Maximum Entropy models for DCA, which differ in the choice of fitted observables. The usual Potts model, in which all first and second moments of the seed MSA are fitted, is labeled as *potts*, while we also consider a “pseudo” Hidden Markov Model (*phmm*), with Hamiltonian

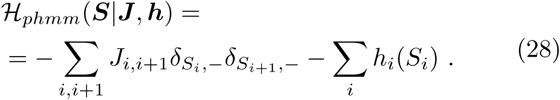

The *phmm* can be thought of as a profile model playing the role of the emission probabilities of a Hidden Markov Model (HMM), plus a pairwise interactions *J*_*i,i*+1_ between neighboring gaps. This interaction is related, in our mapping to a HMM, to the probability of switching, between two consecutive positions, from a “gap” to a “match” state and vice-versa. We also considered other variations of the Potts model, such as a model in which we do not fit the second moment statistics of non-neighboring gaps (i.e. long-range gap-gap couplings are set to zero). The motivation behind this choice is that, if DCA couplings are interpreted as predictors for the (conserved) three dimensional structure, gapped states do not carry any information about co-evolution of far-away positions. However, we found that these other variations do not bring additional insight with respect to the *potts* and *phmm* models, so we restrict here the presentation to these two choices.

All these models are learned on a seed alignment using a standard Boltzmann machine DCA learning algorithm [47]. We used a constant learning rate of 5·10^−2^ for most protein families, and 10^−2^ for all RNA families and for the longest protein families we used. Because the seed often contains very few sequences, we need to introduce a small pseudo-count of 10^−5^ to take into account non-observed empirical second moments. The Boltzmann machine performs a Monte Carlo sampling of the model using 1000 independent chains and sampling 50 points for each chain (in total the statistics is thus computed using 5 10^4^ samples). Equilibration and auto-correlation tests are performed to increase or decrease, if needed, the equilibration or the sampling time of the Monte Carlo.

It is important to keep in mind that the models inferred from the seeds are “non-generative” because of the reduced number of sequences (samples generated from these models, due to a strong over-fitting, are extremely close to seed sequences), but nonetheless they are accurate enough to be used as proper cost functions for our alignment tool. We also mention that models inferred from Pseudo-Likelihood maximization [21, 49], which are also known to be non-generative [47], can be equivalently used for the alignment method described in this work.

### B. Insertion penalties

We determine the parameters of the affine insertion penalties using the statistics of the insertions of the seed alignment. Recall that the number of insertions between positions *i* and *i* − 1 is Δ*n* = *n*_*i*_ − *n*_*i*−1_ − 1, as illustrated in Fig. 3. Motivated by the empirical statistics of insertions in true seeds, we model the probability of Δ*n* as

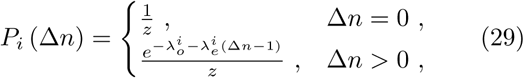

where

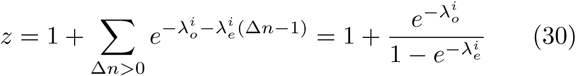

is the normalization constant and 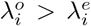 are the costs associated with the opening and the extension of an insertion as in the score function defined in Eq. (3). Because the learning of the parameters is done independently for each position *i*, for the sake of simplicity we will drop the index *i* in the following.

**FIG. 3.**
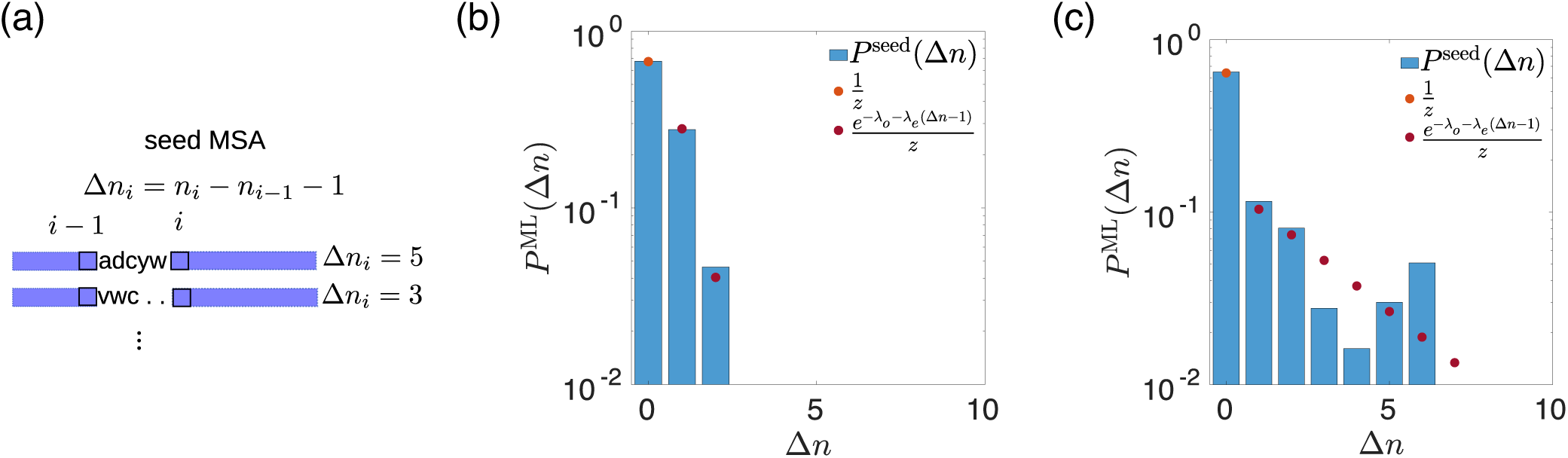
Inference of Insertion penalties. (a) Schematic representation of the Δ*n* variables: the number of insertions between two consecutive positions in the alignment can be computed from the pointer variables ***n***. (b),(c) Examples of fitting of the empirical probability of Δ*n* using our maximum likelihood approach for (b) position 25 and (c) position 92 for RF00162. In (c) the data distribution does not show an exponential profile but our approximation fits well the empirical probability for most of the observed Δ*n*.

We determine the values of *λ*_*o*_, *λ*_*e*_ by maximizing the likelihood 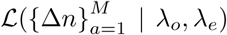 of the data, i.e. the *M* sequences of the seed, given the parameters, and adding L2-regularization terms in order to avoid infinite or undetermined parameters. Imposing the zero-gradient condition on the likelihood leads to a closed set of non-linear equations for the maximum likelihood estimators, given in the SI. These equations can be solved, for example, by a gradient ascent scheme in which we iteratively update

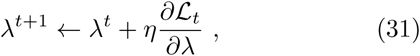

for both *λ*_*o*_ and *λ*_*e*_, until numerical convergence (more precisely, when the absolute value of the gradient is less then 10^−4^). The learning rate is *η* = 10^−3^. Note that the empirical distribution can differ from that of our model: for instance, we often encounter positions where either no insertion is present within the seed or the distribution of the positive Δ*n* is not exponential. In the first case, our maximum likelihood approach cannot be applied as it is: in order to apply it we pretend that the probability of observing at least one insertion is equal to a small parameter *E* so that we can slightly modify *P*^seed^(Δ*n* = 0) = 1−*ϵ*. In our work this parameter has been set to *ϵ* = 10^−3^. In the second case we notice that the distribution given by the fit is anyway a nice approximation of the true one. Some examples are reported in Fig. 3.

### C. Gap penalties

The gap state is treated in DCA models as an additional amino acid but, by construction of the MSA, it is actually an *ad hoc* symbol used to fill the vacant positions between well-aligned amino acids that are close in the full-length sequence ***A*** and should be more distant in the aligned sequence ***S***. Thus the proper number of gaps for each candidate alignment is often sequence dependent and not family dependent: the one-point and two-points statistics of gaps computed from the seed may not be representative of the full alignment statistics. Yet, the couplings and the fields of the DCA models learned from the seed tend to place gaps in the positions mostly occupied by gaps in the seed. This may lead to some bias depending on the seed construction: we notice that if we create seeds using randomly chosen subsets of Pfam [12] alignments produced by HMMer [11], our alignment method, DCAlign, is likely to produce very gapped sequences. In these cases gaps appear very often, more often than any other amino acid, indicating that our cost function encourages the presence of gaps. Real seeds are instead manually curated and therefore they generally contain few gaps. Even though the Potts models learned from this kind of seeds are less biased, we anyway need to check whether the issue exists and, if needed, treat it. To do so, our idea is to introduce additional penalties to gap states, *µ*_ext_ and *µ*_int_, as we discussed in Sec. II B in the definition of the cost function. Notice that the distinction between “internal” and “external” gap penalties allows us to differentiate between gaps that are artificially introduced (as in the case of the internal ones) and gaps that reflect the presence of well aligned but shorter domains or fragments, of effective length *L*frag *< L*. In this last case some “external” gaps are needed to fill the *L*− *L*_frag_ positions at the beginning or at the end of the aligned sequence.

Contrarily to the insertions penalties, the gap penalties cannot be directly learned from the seed alignment via an unsupervised training (as their statistics is already included in the Potts model to begin with), but they can be learned in a supervised way. A straightforward procedure consists in re-aligning the seed sequences using the insertions penalties and the DCA models (*potts* or *phmm*) described in Sec. IV A for several values of *µ*_ext_ and *µ*_int_. The best values of the gap parameters are those that minimize the average Hamming distance between the re-aligned seed and the original seed sequences. We performed this supervised learning by setting the values of *µ*_ext_ *∈* [0.00, 4.00] and *µ*_int_ *∈* [0.00, 4.00]. These intervals have been chosen after several tests in a larger range of variability, also including negative values (that favor gaps) and very large values compared to the typical parameters of the Potts models. We observed that: (i) favoring gaps is always counterproductive, and (ii) there exists a threshold, usually around 4, beyond which no gap is allowed in the sequence, which is also counter-productive. For these reasons, and because of the high computational effort required to re-align the sequences several times, we decided to use the interval [0.00, 4.00] with sensitivity 0.50 leading to 81 re-alignments of the seed sequences.

This method works for seeds that contain a large number of sequences (typically *M >* 10^3^) but it fails completely when dealing with “small” seeds. In this case, whatever the value of (*µ*_ext_, *µ*_int_), the re-aligned sequences are always identical to the original ones, resulting in an average Hamming distance equal to zero. Indeed, the energy landscape of models learned from few sequences is populated by very isolated and deep local minima centered in the seed sequences. When the algorithm tries to re-align an element of the training set, it is able to perfectly minimize the local energy and re-align the sequence with no error whatever the additional gap penalty. For short seeds, instead of realigning the seed, we thus extract 1500 sequences from the full set of unaligned sequences (a *validation* set), which we align by varying the gap penalties, always in the range [0.00, 4.00]. We call 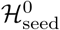 the DCA Hamiltonian inferred on the seed, and we infer new Hamiltonians 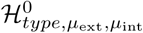 (with *type ∈ {phmm, potts,*}) on all the multiple sequence alignments of the validation set. We then choose the best parameters *µ*_ext_ and *µ*_int_ as those that minimize the symmetric Kullback-Leibler distance between 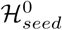 and 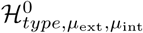 (the precise definition is given in section V). In other words, we select the best gap penalties as those that produce a validation MSA as statistically close as possible to the seed alignment.

We underline that the values of the penalty parameters also depend on the choice of the gauge for the DCA parameters: in fact, the advanced mean-field equations are not gauge invariant and depending on the choice of the gauge we can have different (optimal) values for the gap penalties. All results shown in this work use the zero-sum gauge for the DCA parameters.

## V. EXPERIMENTAL SETUP

### A. Pipeline

The experimental setting we propose here is the same adopted by state-of-the-art alignment softwares such as HMMer [50], and Infernal [14]. From the seed alignment we learn all the parameters that characterize our score function and we apply our alignment tool to all the un-aligned sequences that contain a domain compatible with the chosen family. More precisely, once a seed is selected (we used the hmmbuild -O function of the package HMMer to obtain the aligned seed), we learn the model, the insertions parameters and the gap penalties as described in Sec. IV. A scheme of our training method is shown in Fig. 4.

**FIG. 4.**
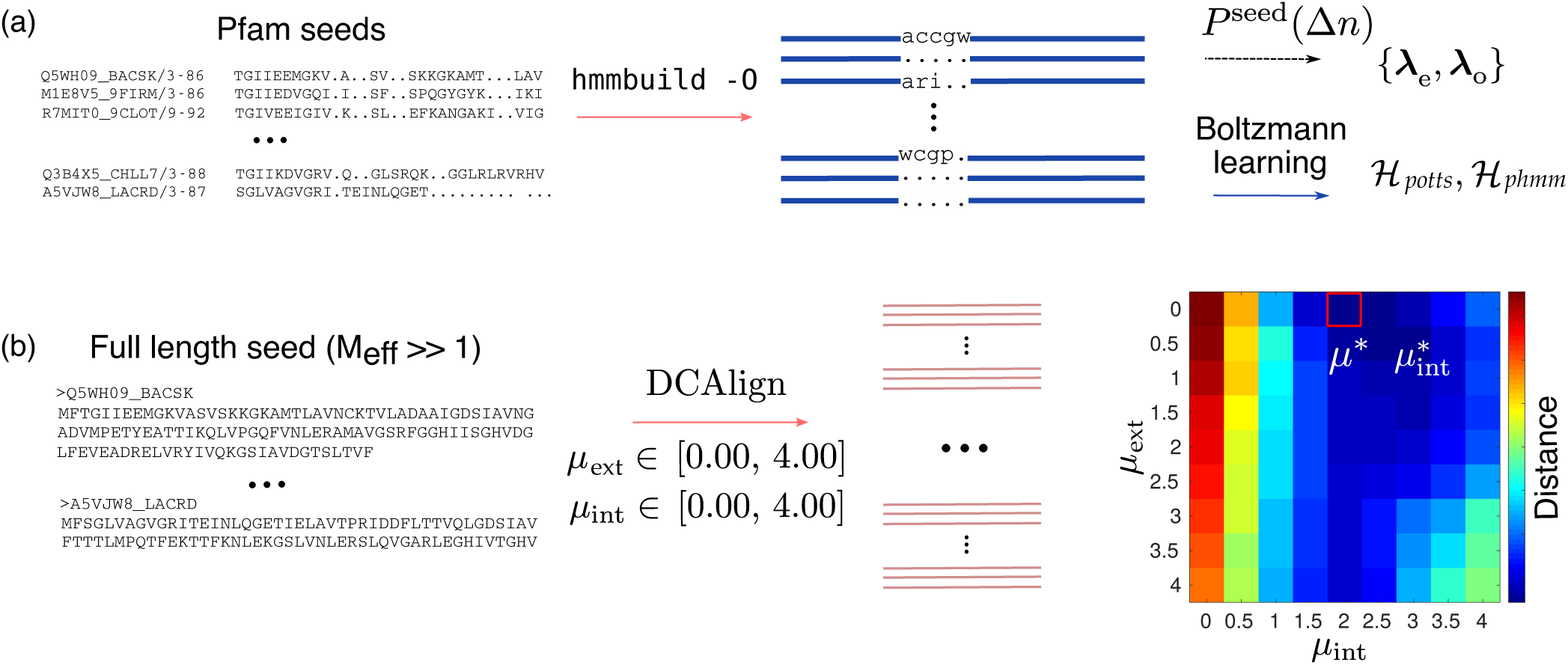
Scheme of the training process for DCAlign. In (a) we show the first step of the learning. We build the aligned seed of a Pfam family using `hmmbuild -O` to detect the matched amino-acids (blue line) and the insertions (shown in lower case letters and “.”). From these data we learn the DCA Hamiltonian and the affine insertion penalties. In (b) we pictorially describe the learning of the gap penalties. Here we take into account the seed itself (not only the aligned part but the entire sequences) and we try to align it using the parameters inferred in step (a) using all the 81 possible combinations of *µ*_ext_ and *µ*_int_, each spanning the interval [0.00, 4.00]. We compare each candidate alignment to the seed alignment, directly using the Hamming distance. The best set of parameters is that minimizing this metric. The plot in (b) shows the (average) Hamming distance of the true and re-aligned seed for the PF00677 family.

After training, our cost function is fully determined. Unaligned sequences are then taken from the full length sequences of the corresponding family in the Pfam database [12] (we do not face here the problem of detecting homologous sequences). Note that, like HMMer, we also include the seed sequences in the sequences to be aligned, in order to obtain a more homogeneous MSA and test the quality of the re-alignment of the seed. We do not consider the entire sequences, whose length *N* is often much larger than *L*, but a “neighborhood” of the hit selected by HMMer. In practice we add 20 amino-acids at the beginning and at the end of the hit resulting in a final length *N* = 20 + *L* + 20. We performed the same experiment using *N* = 50 + *L* + 50 for PF00684 and the resulting sequences were identical to those obtained from a shorter hit. The method seems to be stable for reasonable values of *N*, i.e. *N* ∼ *L*. For RNA sequences, this pre-processing is not needed, because the full length sequences downloaded from Rfam already have a reasonable length. For most families, we have aligned all the full length sequences (the size of the test sets is specified in Table II) and only in few cases, for particularly large families, we uniformly pick at random *N*_seq_ = 10^4^ sequences to align.

**TABLE II.** Features of the protein and RNA families used in this work. We show here the length *L* of the sequences for each family, the value of *M* and *M*_eff_ [20] for the seed alignment, the number of PDBs used to determine the true contact maps based of real observations of the domains structure, the number of the sequences, *N*_seq_, to be aligned by our methods and the value of the gap penalties associated with each family and Hamiltonian.

| Identifier | $L$ | $M$ seed | $M_{\text{eff}}$ seed | PDB | $N_{\text{seq}}$ | $\mu_{\text{ext}}$ | | $\mu_{\text{int}}$ | |
| --- | --- | --- | --- | --- | --- | --- | --- | --- | --- |
| | | | | | | $\mathcal{H}_{\text{potts}}$ | $\mathcal{H}_{\text{phmm}}$ | $\mathcal{H}_{\text{potts}}$ | $\mathcal{H}_{\text{phmm}}$ |
| PF00035 | 67 | 81 | 81 | 73 | 19751 | 2.50 | 2.00 | 0.00 | 2.00 |
| PF00677 | 87 | 1878 | 1518 | 9 | 14683 | 0.00 | 0.50 | 2.00 | 2.00 |
| PF00684 | 67 | 1512 | 1349 | 3 | 10000 | 0.00 | 0.00 | 2.50 | 2.00 |
| PF00763 | 116 | 1389 | 1355 | 24 | 10000 | 1.50 | 2.50 | 1.00 | 1.50 |
| RF00162 | 108 | 433 | 241 | 25 | 6026 | 3.50 | 3.50 | 3.00 | 4.00 |
| RF00167 | 102 | 133 | 105 | 49 | 2631 | 0.50 | 1.50 | 2.00 | 4.00 |
| RF01734 | 63 | 287 | 287 | 6 | 2017 | 1.00 | 0.00 | 2.00 | 1.50 |
| RF00059 | 105 | 109 | 83 | 24 | 12558 | 0.00 | 0.50 | 1.50 | 1.50 |

We then apply DCAlign using the approximations we discussed in Sec. III, namely the finite temperature and zero-temperature MF method, to each candidate sequence and we add to our MSA the aligned sub-sequence that has the lower energy (insertion and gap penalties excluded). Whenever our algorithms do not converge to an assignment of the variables that satisfies all the hard constraints, we apply the “nucleation” procedure explained in Sec. III C that, by means of the approximated marginal probabilities, gives rise to feasible alignments.

### B. Observables

To assess the quality of the MSAs generated by DCAlign (that differ in the score function being used to align) and to compare them to the state-of-the-art alignments provided by HMMer (or Infernal), we consider the following observables.

- *Sequence-based metrics*. When comparing two candidate MSAs of the same set of sequences (a “reference” and a “target”), it is possible to compute several sequence-wise measures such as the following metrics (normalized by *L*, the length of the sequences):
  – the Hamming distance between the two alignments of the same sequence in the reference and target MSAs;
  – Gap_+_: the number of match states in the aligned sequence of the reference MSA that have been replaced by a gap in the target MSA;
  – Gap_−_: the number of gaps in the aligned sequence of the reference MSA that have been replaced by match states in the target MSA;
  – Mismatch: the number of amino-acid mismatches, that is the number of times we have match states in both sequences, reference and target, but to different amino acids (positions) in the full sequence ***A***.
- *Proximity measure*. Consider two different MSAs of the same protein or RNA family. We can compute, for each candidate sequence 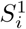 of the first set, the Hamming distance *d*_*H*_ with respect to all the sequences of the second set. We then collect the minimum attained value, i.e.

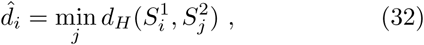

which gives the distance to the closest sequence in the other MSA. The distribution of 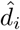, or some statistical quantity computed from them (such as the average or the median value) provides a good measure of “proximity” between the two sets. We will show below a few examples using the full alignment of a protein family as a first set, and the seed sequences as the second one.
- *Symmetric Kullback-Leibler distance*. Another convenient global measure is the symmetric Kullback-Leibler distance between a Boltzmann equilibrium model learned from the seed alignment (the “seed” model) and another model learned from a candidate MSA (the “test” model). In general the symmetric Kullback-Leibler divergence is a measure of “distance” between two probability distributions and it is defined, for arbitrary densities *𝒫*_1_(***x***) and *𝒫*_2_(***x***) of the variables ***x***, as

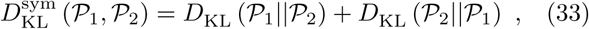

where *D*_KL_ is the Kullback-Leibler divergence

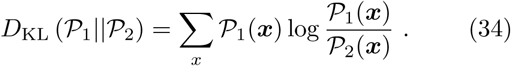 In our context, the symmetric KL distance can be efficiently computed through averages of energy differences as

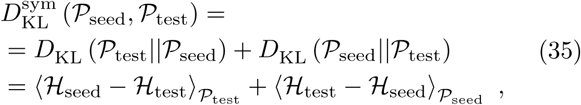

where *𝒫*_(.)_ is the Boltzmann distribution associated with the energy ℋ_(.)_, and the brackets ⟨…⟩_*P*_ denote the expectations with respect to *𝒫*, which can be easily estimated using a Monte Carlo sampling (in contrast to the normal *D*_KL_, which depends on the intractable normalization constants, i.e. the partition functions, of the densities *𝒫*_(.)_). We run the comparison using two different models for ℋ, that is a Potts model and a profile model.
- *Contact map*. The couplings of DCA models can be used to detect the presence of physical interactions between pairs of sites, which are distant in the one-dimensional chain but in close contact in the three-dimensional structure. A good score that indicates a direct contact is the (average-product corrected) Frobenius norm of the coupling matrices, defined as

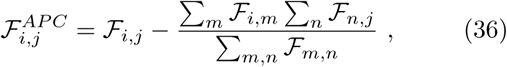

where

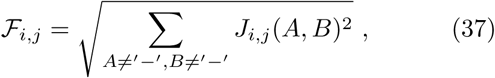

and the couplings are in the zero-sum gauge. We thus compare predicted contact maps obtained using the parameters learned from our alignments and those obtained by HMMer. For this purpose, we use the PlmDCA method to learn the couplings from the alignments, because it is faster than the Boltzmann machine and it is known to be reliable for contact prediction [21]. The ground-truths denoting the physical interactions in each protein are obtained running the *Pfam interactions* package [51]. Two sites are said to be in contact if the minimum atomic distance among all the atoms and among all the available protein structures is less than 8 Å.

## VI. TEST ON SYNTHETIC DATA

Here we describe two experiments on synthetic data, constructed to compare the performances of DCAlign to state-of-the-art methods in extreme settings: one dataset presents conserved but not coevolving sites (i.e. strong variations in amino-acid frequency on each site but no correlation between distinct sites), while the other presents not conserved but coevolving sites (i.e. uniform frequency 1*/q* of amino-acids on each site, but strong correlations between distinct sites).

### A. Conservation

The first MSA is generated from a non-trivial profile model, in which the empirical probability of observing any of the possible amino acids is position-dependent and it is not uniformly distributed among the possible states. The generative model used in this case is the profile model of the PF00018 family, which can be easily learned from the empirical single site frequencies. From this model we generate 5·10^4^ “aligned” sequences (the “ground-truth”), to which we randomly add some insertions, according to the affine insertion penalty distributions learned from the PF00018 seed (Sec. IV B), and 20 uniformly randomly chosen symbols at the beginning and at the end of the aligned sequence. We split this alignment into a training set of 2.5·10^4^ sequences, which we use as seed alignment to learn the insertion and gap penalties (using the scheme for abundant seeds) and a Potts model. We align the remaining 2.5·10^4^ sequences, used as test set. For comparison, we build a Hidden Markov Model using hmmbuild of the HMMer package on the training set and we align the test sequences through the `hmmsearch` tool.

We show in Fig. 5(a) the histograms of the (normalized) Hamming distances, Gap_+_, Gap_−_ and Mismatches of the MSA obtained by DCAlign and HMMer compared to the ground-truth. We observe that DCAlign is able to find the correct hits and to align them in a more precise way if compared to HMMer. In fact, the Hamming distance distribution is shifted to smaller values, suggesting that the number of errors, per sequence, is smaller than that obtained by HMMer. The nature of the mistakes seems to be linked to the presence of mismatches in the case of HMMer, while DCAlign (less often) equally likely inserts more or less gaps, or a match to the wrong symbol. While DCAlign is in principle constructed to exploit covariation, these results show that even in cases in which, by construction, there is no covariation signal, DCAlign is able to perform equally good (or even better) than state-of-the-art methods.

**FIG. 5.**
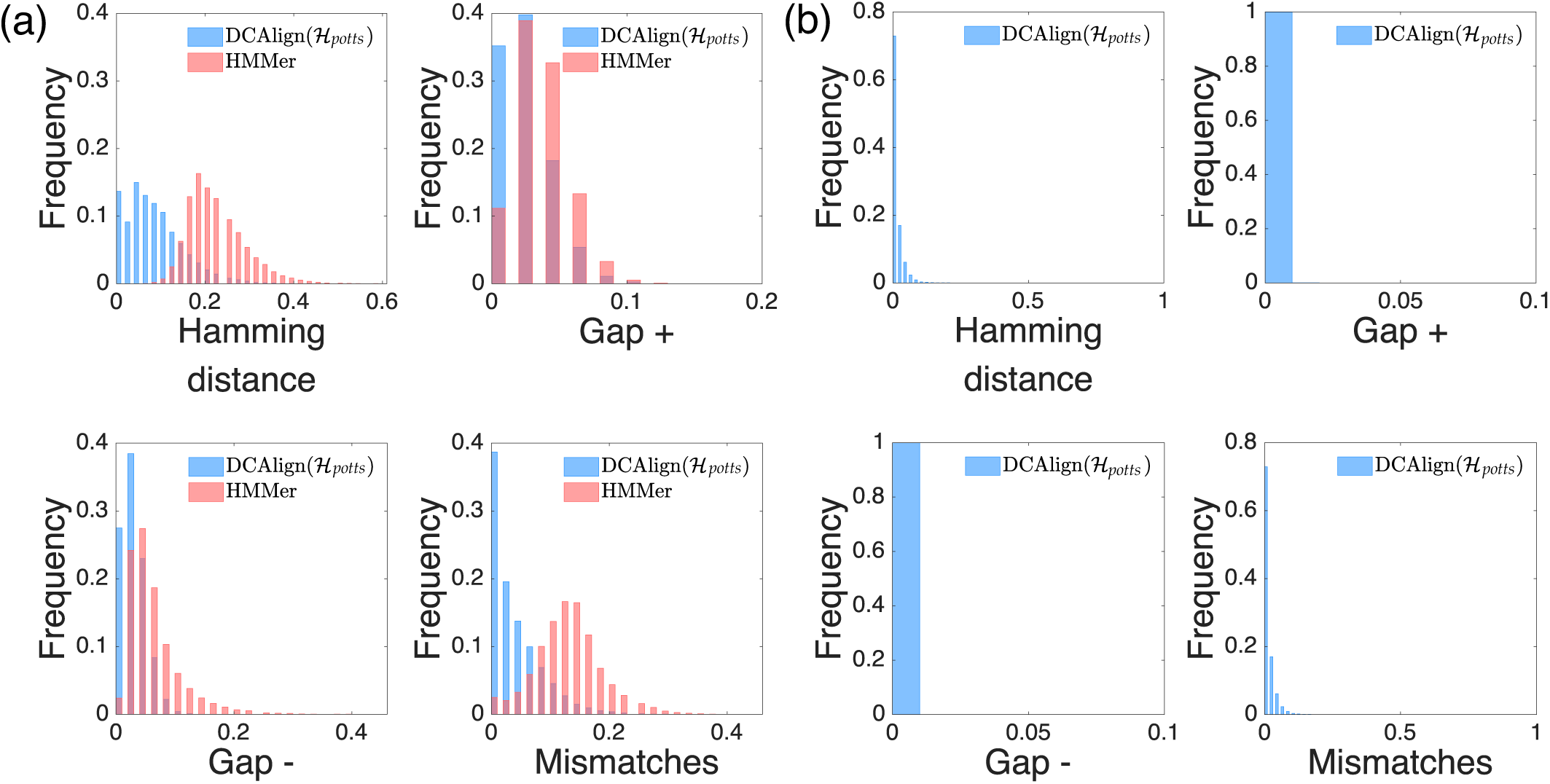
Comparison between DCAlign and HMMer for synthetic data. Panels (a) and (b) show the histograms of the normalized metrics (Hamming distance, Gap_+_, Gap_−_ and Mismatches), respectively, in the case of conserved sited only, and of correlated pairwise columns only. Here, the reference is the ground truth and the target is the alignment of DCAlign (blue) or HMMer (red). In (b) HMMer results are not shown because `hmmsearch` does not find any relevant hit.

### B. Covariance

The second experiment is instead focused on correlated data. We ad-hoc construct an alignment whose first moment statistics resembles that of a uniform distribution, i.e. the probability of observing any amino-acid, in any column of the seed, is 1*/q*. In other words, we construct the data in such a way that no conserved sites are present. At the same time, we force the sequences to show coevolving (i.e. correlated) sites, such that the empirical probability of observing a pair of amino acids is different from that o<u>btained in</u> the uniform distribution, i.e. 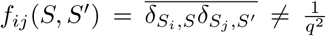 for some (*i, j*), where the overline indicates the empirical average. To construct a dataset with this statistics, we use as generative model a Potts model with 4 colors (like the RNA alphabet, without the gap state) having non-zero couplings 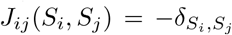 (i.e. an anti-ferromagnetic Potts model) and no fields. The non-zero couplings are associated with the links of a random regular graph of 50 nodes and degree 5. The presence of the links ensures the appearance of non-trivial second moments while, in order to avoid “polarized” sites, we sample the model (i.e. the Boltzmann distribution associated with this Hamiltonian) at temperature 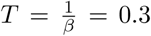, which is deep in the paramagnetic phase of this model [52]. We perform the same training pipeline presented in Sec. VI A except for the learning of the gap penalties: because there are no gap states, we set *µ*_ext_ = 0 and *µ*_int_ = 0.

In this case, due to the absence of any conservation, `hmmsearch` does not find any eligible hit. In fact, HMMer tries to align the sequences via a computationally exact recursion on a HMM, but it has no information to exploit while setting up the HMM from the training set, because all amino-acids are equally likely to occur in each column. This represents, of course, the worst-case scenario for HMM-based methods. On the contrary, the couplings of the learned Potts model allow DCAlign to align this kind of sequences. We remark that contrarily to HMMer, DCAlign has complete information on the statistics of the training alignment, up to second order covariances. However, being a heuristic method, it sometimes fails to achieve the (global) minimum of the cost function and converges to a local minimum, which depends on the initialization of the target marginals. Re-iterating the MF equations using 10 different seeds of the random number generator suffices to reach the proper minimum at least once, for the majority of sequences. We remark that this issue is present only when the MSA does not show any conserved site and thus the algorithm has no easy “anchoring” point, which surely helps lifting the degeneracies in the alignment procedure. For protein and RNA families presented below the algorithm seems stable and only one minimum emerges upon re-initialization of the marginal probabilities of the algorithm. We quantitatively measure the performance of DCAlign using four sequence-based metrics and the energies associated with the aligned sequences. For this experiment, we refer to the output of our algorithm as the aligned sequence that has the minimum energy among the 10 trial re-iterations of the algorithm. We report the distance metrics in Fig. 5(b): the distribution of the Hamming distances suggests that DCAlign can almost perfectly align the majority of the target sequences. Indeed, as shown in Fig. 6, the energies (the Potts Hamiltonian alone or the full cost function which includes the gap and insertion penalties) are identical or very close to the energies of the true sequences. Only 0.08% of the aligned sequences have a Hamming distance density larger than 0.30 (i.e. 15 missed positions over 50). This fraction is so low to be invisible in the histograms of Fig. 5(b). These extreme cases, in which our algorithm converged to a local minimum in every trial we performed, are represented as purple points in the scatter plots in Fig. 6.

**FIG. 6.**
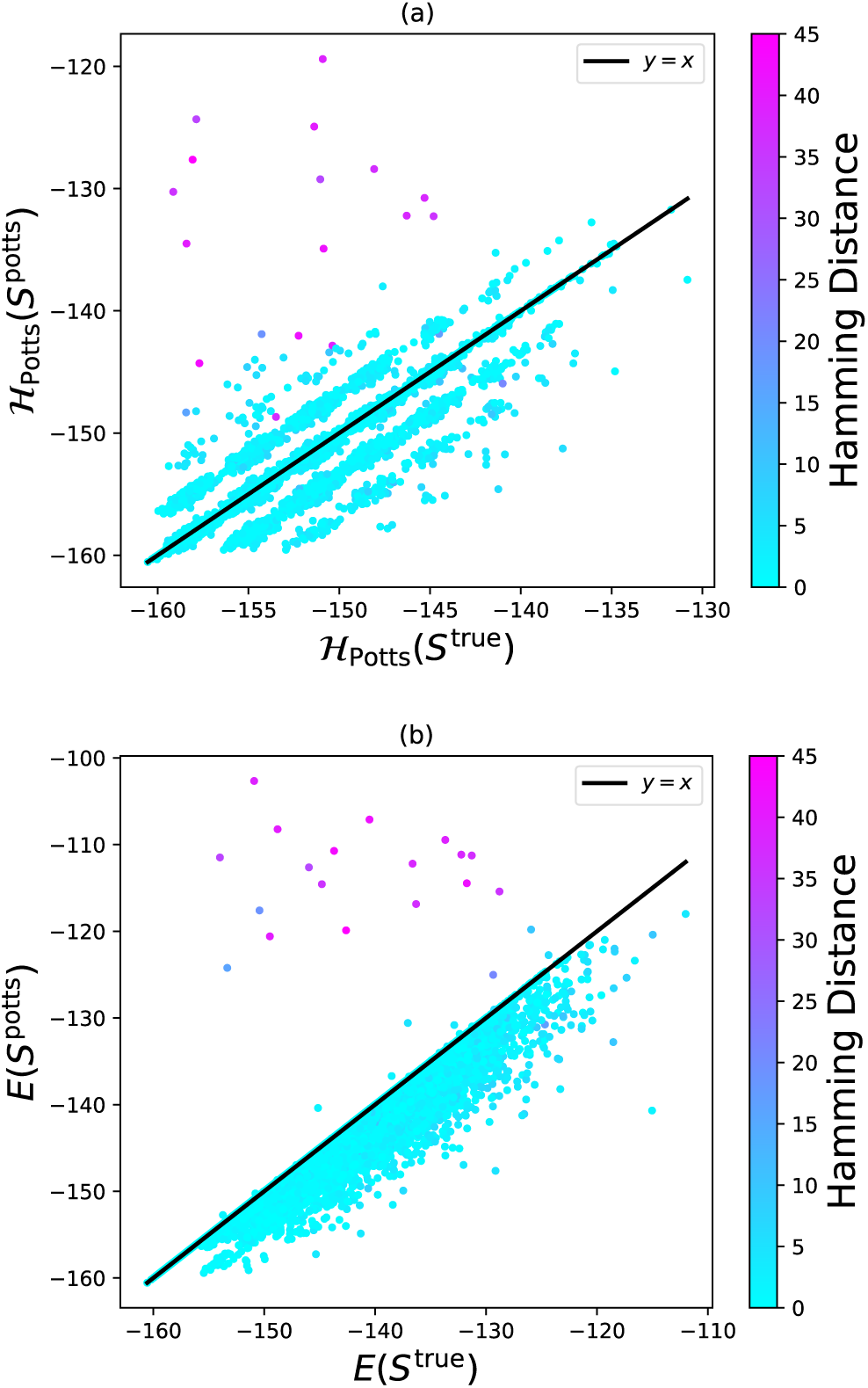
Energies of synthetic sequences. We show here the scatter plots of the DCA energy (or Potts Hamiltonian) in (a) for the true synthetic sequences (x-axis) against the ones aligned by DCAlign(ℋ_*potts*_) (y-axis). In (b), same plot using the total cost function *E*.

## VII. TEST ON PROTEIN AND RNA FAMILIES

### A. Choice of families

We show here the performance of our alignment method for several RNA and protein families. In particular we select the families PF00035, PF00677, PF00684, PF00763 from the Pfam database (https:://pfam.xfam.org release 32.0) [12], and RF00059, RF00162, RF00167 and RF01734 from the Rfam database (https:://rfam.xfam.org release 14.2) [53]. The number of sequences, length of the models, and gap penalties used in the simulations are reported in Table II.

We restrict our analysis to “short” families, having *L* of at most 100 positions, in order to avoid a significant slowing down of the alignment process. The seed of PF00035 contains very few sequences, contrarily to PF00677, PF00684, PF00763, which have been chosen because of their large effective number of seed sequences *M*_eff_ *>* 1000 (after a standard re-weighting of close-by sequences [20]). We thus always infer the gap penalties according to the abundant seed protocol, except for PF00035. As reference for comparison, we consider the alignments produced by HMMer [50] and already available in the Pfam database. We also perform the alignment of several RNA families, for which secondary-structure knowledge is necessary to obtain good alignments with standard tools. We compare our estimate against that obtained by the state-of-the-art package Infernal [14] which, indeed, employs the secondary structure of the target domains in order to build the so-called covariance model used to align. Note that, on the contrary, DCAlign does not use any secondary structure information in the training procedure (but DCA is able to predict the latter [54]). As a further comparison we also learn a Hidden Markov Model (using `hmmbuild`) and we apply `hmmalign` to the full length RNA sequences. We choose precisely these families because of their reasonable length, the abundance of the seed sequences, and the large number of available crystal structures, which are useful for the contact map comparison.

### B. Comparison with state-of-the-art methods

As a first comparison, we compute the sequence-based metrics presented in Sec. V B comparing our full alignment to that achieved by HMMer, for protein sequences, or by Infernal, when dealing with RNA families. We show the results for PF00035 in Fig. 7(a-d) and for RF00167 in Fig. 7(e-f), which are representative of the typical scenario for protein and RNA sequences. The distribution of all metrics is mostly concentrated in the first bins (the bin width is set here to 0.01) and decays smoothly at larger distances. The peak in the first bin is more prominent when the Hamiltonian used for the alignment is *phmm* indicating that the sequences aligned by this method are closer to those obtained by HMMer (or Infernal) than the outcomes of DCAlign-*potts*, as one would expect from the similarity between the two alignment strategies (see Sec. IV A).

**FIG. 7.**
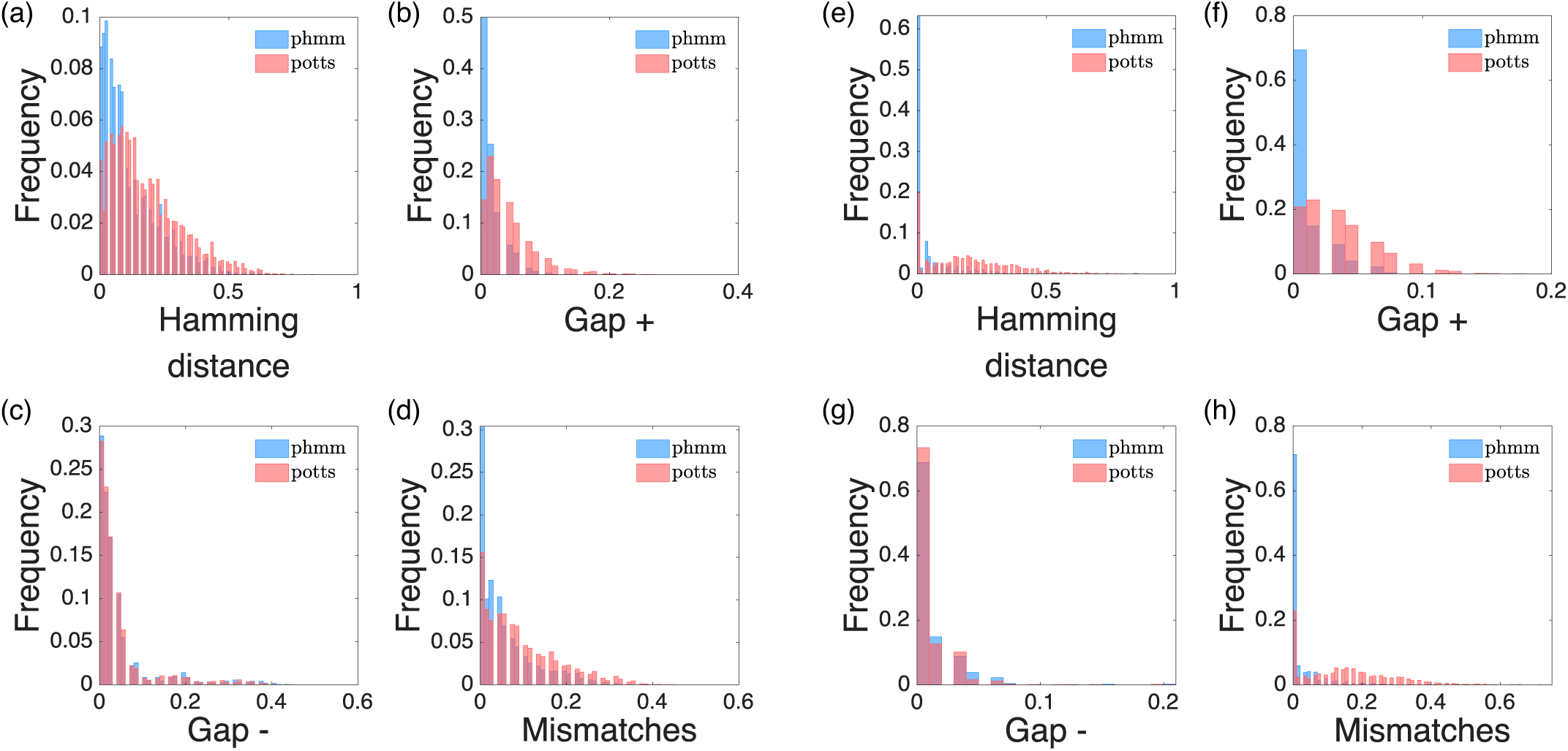
DCAlign vs state-of-the-art methods for PF00035 and RF00167. We plot here the histograms of the Hamming distances, Gap_+_, Gap_−_ and Mismatches for the protein family PF00035 (a, b, c, d) using as reference the HMMer results and as target the DCAlign results, and for the RNA family RF00167 (e, f, g, h) using as reference the Infernal results and as target the DCAlign results.

A notably different behavior emerges for the sequences of the PF00677 family, as shown in Fig. 8(a-d). It is clear from Fig. 8(a) that a large fraction of the sequences aligned by DCAlign differs from those aligned by HMMer by about 40% of the symbols when using ℋ_*potts*_ (the percentage is reduced to about 30% when using ℋ_*phmm*_). The reason seems to be partially linked to the presence of mismatches and to a non-negligible fraction of additional gaps, as indicated by Gap_+_. We notice that, differently from the other families, the seed of PF00677 is composed by several clusters of sequences mostly differing in the gap composition: a copious fraction of them have generally few gaps while some other show two long and localized stretches of gaps. The structure of the seed can be better characterized in the principal components space, as depicted in Fig. 8(e), where we plot the projections of the seed sequences in the space of the first two principal components as filled circles. The colors refer to the density of sequences in the (discretized) space. To understand the disagreement between HMMer and our methods we superimpose the projections of the sequences responsible for the huge peaks in the Gap_+_ histogram in the principal component space of the seed sequences, using brown crosses for the sequences aligned by HMMer and pink diamonds for those found by DCAlign-*potts* (note that none of these sequences is a re-alignment of a seed sequence). Only a small fraction of the HMMer sequences overlap with the central and poorly populated cluster while the projections obtained from sequences aligned by DCAlign-*potts* lie on a well defined and populated cluster. We thus conclude that looking at the gap composition of sequences is not sufficient in this case to understand the different behavior of HMMer and DCAlign. A more accurate analysis in the principal components space suggests that the sequences obtained by HMMer are probably miscategorized, at variance with DCAlign sequences that are in agreement with the seed structure.

**FIG. 8.**
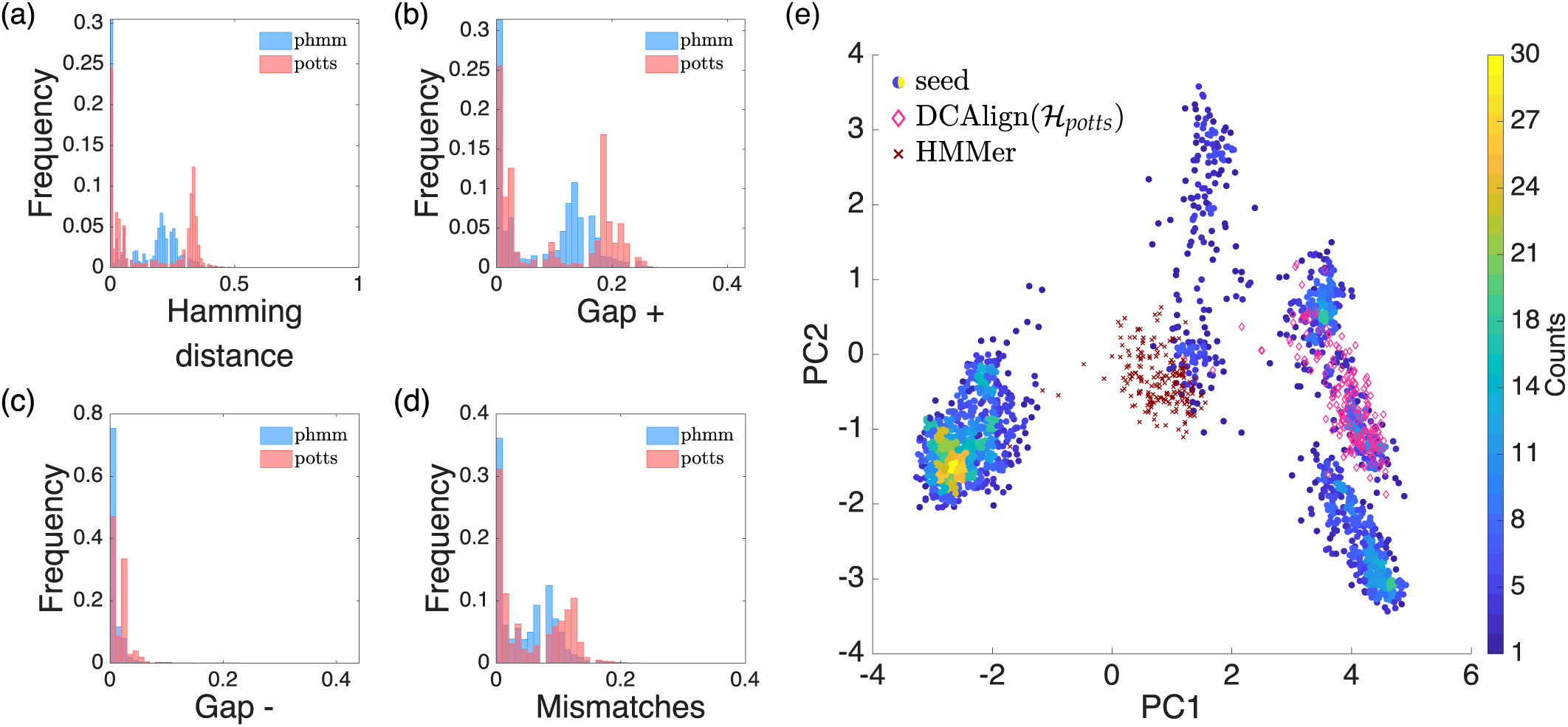
DCAlign vs HMMer for PF00677. In (a, b, c, d) we plot the Hamming distance, Gap_+_, Gap_−_ and Mismatches using the sequences aligned by HMMer as reference and those obtained by DCAlign as target. In (e) we plot the projections of the seed sequences in the first two principal components of the seed space; the color scale denotes the density of the space. The additional sequences (depicted as pink diamonds if aligned by DCAlign-*potts* or as brown crosses if aligned by HMMer) are responsible for the red peak around 0.2 in panel (b).

In summary, although for most of the families analyzed here (the distribution of the four metrics for the remaining families are shown in the SI) the sequences aligned by DCAlign are very similar to those obtained by HMMer or Infernal, the PF00677 family suggests a different scenario, in which DCAlign is able to learn some non-trivial seed features and to align the target sequences according to them.

### C. Comparison with the seed

In this section, we compare the statistical properties of the MSAs obtained by DCAlign with those of the seed.

#### 1. Kullback-Leibler distances

The statistics of a MSA can be characterized in terms of a statistical (DCA) model. Depending on the complexity of the model, a certain set of observables are fitted from the MSA. For instance, in a profile model only the first moments are fitted, while in a Potts model we can also fit the information about second moments. These statistical models define a probability measure over the space of sequences and thus characterize a given protein/RNA family. We consider here the seed sequences as our ground truth, and we thus consider that a model learned from the seed is the one that better characterizes the protein/RNA family under investigation. We then infer a second model from the full set of aligned sequences, and we ask how different is this model from that learned from the seed. To answer this question we compute the symmetric Kullback-Leibler divergence 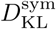 between the two models, see Sec. V B, which must be intended as a statistical measure of distance between the seed and the set of aligned sequences. In order to fairly compare DCAlign, HMMer and Infernal, which by construction treat differently the pairwise co-variation of the MSA sites, we learn, from a seed and from the test alignment, a profile model ℋ^Prof^ and a Potts model ℋ^Potts^.

We show in Fig. 9(a,b) the results for all families and all methods, when the model learned is a Potts model or a profile model, respectively. We notice that the alignments produced by DCAlign-*potts* always, for the Potts case, and very often, for the profile case, minimize 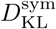 with respect to the seed. Infernal is very effective when dealing with RNA sequences but not as good as DCAlign-*potts* for the majority of the cases. HMMer always produces the largest distance (except for PF00763 where basically all methods perform equally good), in particular for RNA families. We mention that alignments produced by hmmalign present aligned sequences that always show long concatenated gaps at the beginning and at the end of the sequence, differently to the Rfam full alignment, the seed sequences and the outputs of DCAlign. This partially explains the difference with respect to the other alignment tools.

**FIG. 9.**
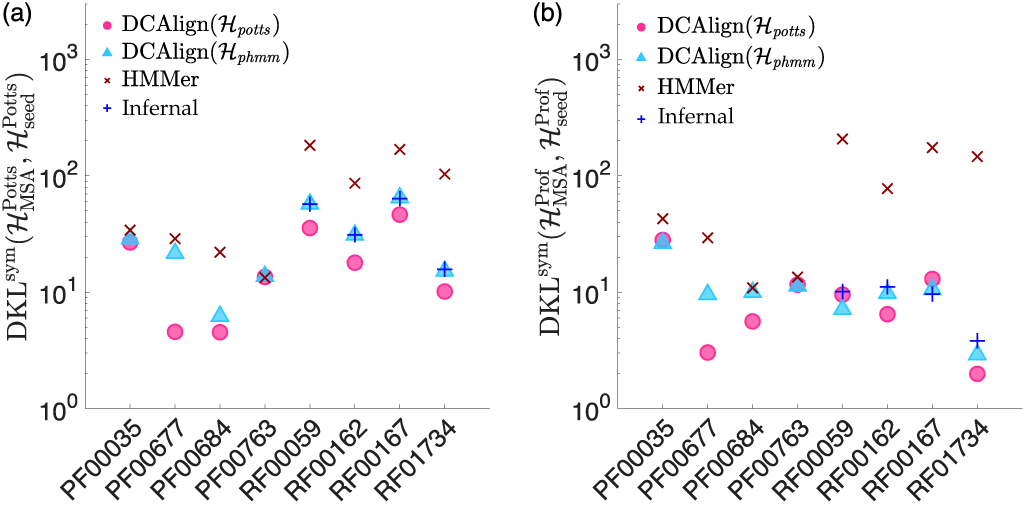
Symmetric Kullback-Leibler distances. We plot the symmetric KL distance between the MSA and the seed alignment, computed via a Potts model (a) or a profile model (b), for all the families and all the alignment methods we considered.

These results suggest that the more we use additional information within the alignment process (in particular, when learning ℋ_*potts*_ we employ all positions and amino-acid dependent pairwise energy function), the closer the final alignment will be to the seed. Surprisingly, this feature is retrieved even when the model learned from the full set of aligned sequences uses less information than the model used to align, e.g. for the profile model used in Fig. 9(b). Of course, there is a tradeoff, because including additional statistical properties of the seed in the alignment process requires a larger seed.

#### 2. Proximity measures

We present here a sequence-based comparison between a candidate alignment, i.e. an alignment obtained by DCAlign-*potts*, DCAlign-*phmm*, or HMMer/Infernal, and the seed that will be considered here as the reference alignment. The metrics we use is the proximity measure introduced in Sec. V B. We show in Fig. 10 the distribution of the minimum distances for a representative subset of the families, i.e. PF00677 in panel (a), PF00684 in (b), RF00162 in (c) and RF01734 in (d). Results for PF00035, PF00762, RF00059 and RF00167 are shown in the SI. We notice that for the majority of the families (protein or RNA) the histograms built from DCAlign-*potts* have a large peak in the first bin (which collects distances from 0 to 0.02), suggesting that there exist more sequences in this alignment which are close to the seed than in any other alignment. A large peak at small distance is also observed for Infernal when dealing with RNA families, as seen from the blue histograms in Fig. 10(c) and (d). The Infernal results overlap quite well with those obtained by DCAlign-*phmm*. The histograms produced by HMMer seem to be shifted to larger Hamming distances, thus reflecting a smaller similarity to the seed than all the other methods. Although DCAlign-*phmm* exploits similar information to that encoded in HMMer, the corresponding alignment surprisingly produces, for most of the studied families, results that are more similar to those obtained by DCAlign-*potts* or Infernal.

**FIG. 10.**
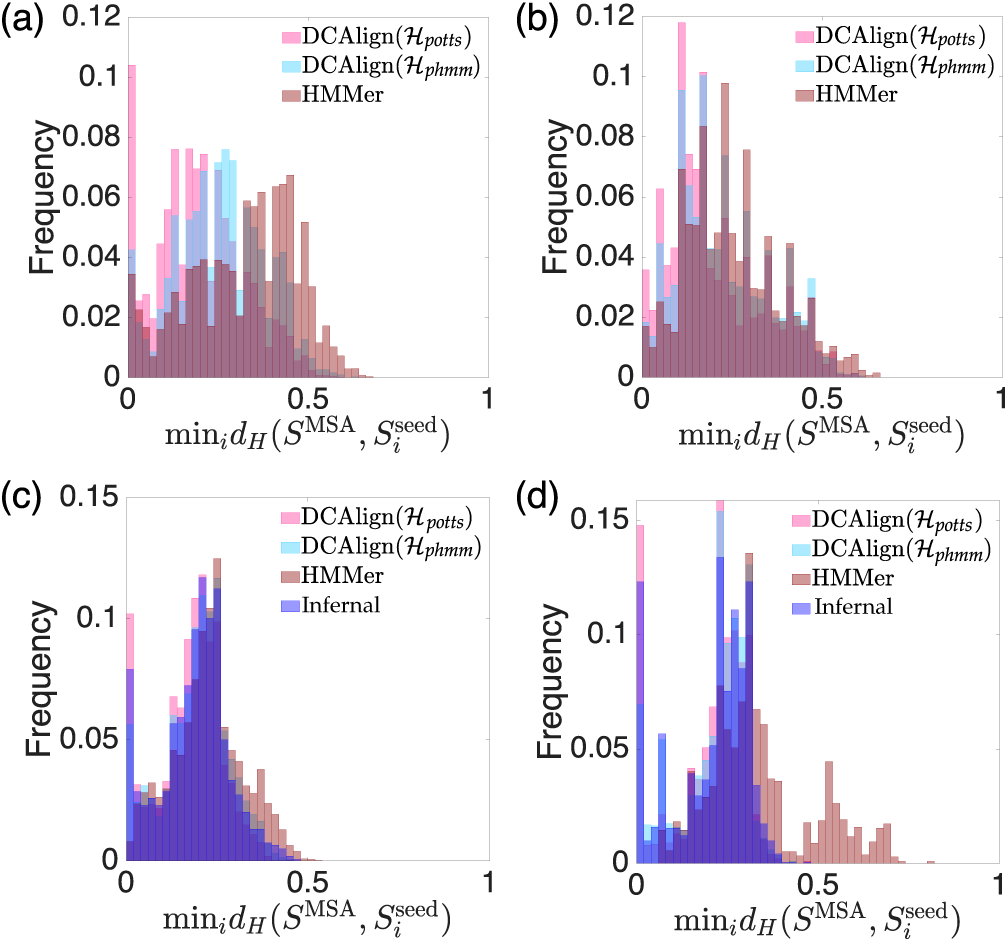
Distribution of proximity measures. Histograms of the minimum distances computed according to Eq. (32) for the full set of aligned sequences obtained by DCAlign-*potts*, DCAlign-*phmm*, HMMer, and Infernal, against the seed. Panels (a), (b), (c) and (d) refer to the families PF00677, PF00684, RF00162, RF01734 respectively.

### D. Contact prediction

An important test of the quality of a MSA is related to the interpretability of the DCA parameters learned from it. As mentioned in Sec. V B, the largest couplings are a proxy for the physical contacts in the folded structure of the protein domains. In Table III we report a summary of the results for three observables associated with the contact prediction: the position of the first false positive in the ranked Frobenius norms, the value of the True Positive Rate (TPR) at 2 *L* and the position at which the TPR is less than 0.80 for the first time. The bold number corresponds to the largest value, and therefore the best performance, among all the methods.

**TABLE III.** Summary of the contact map results. For each protein or RNA family we show here three metrics computed from the PPV curve retrieved from a set of Potts models. ℋ_seed_ is a Potts model learned using the seed sequences alone, while the others are associated with the complete alignments obtained by DCAlign-*potts*, DCAlign-*phmm*, HMMer and Infernal. The chosen observables give the position of the first false positive (First FP), the value of the true positive rate (TPR) computed after 2 ·*L* predictions and the rank at which the value of the true positive rate is smaller than 0.80 for the first time. A perfect prediction is obtained if all the true positive contacts are associated with the highest value of the Frobenius norm, thus the higher the value of these metrics, the better the prediction of the contact maps. We show in bold numbers the best performances, for all metrics and among all the methods.

| Identifier | First FP, TPR(2L), TPR < 0.80 |  |  |  |  |
| --- | --- | --- | --- | --- | --- |
| | $\mathcal{H}_{\text{seed}}^{\text{Potts}}$ | DCAlign ($\mathcal{H}_{\text{potts}}$) | DCAlign ($\mathcal{H}_{\text{phmm}}$) | HMMer | Infernal |
| PF00035 | 9, 0.478, 13 | **35**, 0.754, 98 | 32, 0.799, **134** | 28, **0.791**, 119 | - |
| PF00677 | 22, 0.730, 109 | **47**, 0.759, 147 | 28, 0.747, 128 | 31, **0.793**, **163** | - |
| PF00684 | 20, 0.582, 28 | **29**, 0.672, 101 | 27, **0.694**, **104** | 23, 0.627, 73 | - |
| PF00763 | 80, 0.703, 159 | 89, 0.828, **288** | 85, 0.836, 254 | **106**, **0.849**, 272 | - |
| RF00059 | 18, 0.369, 29 | 29, 0.531, 57 | 29, 0.519, **64** | **37**, 0.519, 59 | 33, **0.566**, **64** |
| RF00162 | 15, 0.306, 52 | 17, 0.449, **69** | **25**, 0.398, 61 | 22, 0.426, 67 | 19, **0.519**, 61 |
| RF00167 | 19, 0.324, 27 | 27, 0.493, 59 | 25, 0.556, 53 | 22, 0.577, **62** | **28**, **0.592**, 57 |
| RF01734 | **10**, 0.300, 16 | **10**, 0.380, 16 | **10**, **0.430**, 16 | **10**, 0.360, 16 | **10**, 0.400, **17** |

We show in Fig. 11 (a,b,c,d) the Positive Predictive Value (PPV) curves (left) and the contact maps (right) for the PF00035, PF00684, PF00763 and RF00162 families respectively (results for PF00677 and the other RNA families are shown in the SI). The PPV curves are constructed by plotting the fraction of true positives TP as a function of the number of predictions (TP and FP), PPV=TP/(TP+FP). The true contact maps are extracted from all the available PDBs and plotted as gray filled squares, while the predicted contact maps are constructed by plotting the Frobenius norms of the DCA couplings that are larger than an arbitrary threshold, here set to 0.20. For RNA sequences the comparison between the predictions and the ground-truth can be performed only using the Frobenius norms associated with the central part of the aligned sequences, because there is no available structural information about the sites on the boundaries. In addition to the predictions obtained from the full set of aligned sequences, we show, for comparison, the predicted contact map obtained from the Potts model inferred from the seed sequences alone. As we can notice from Table III and the plot of the contact maps, there is no strategy that clearly outperforms the others (except the poor results of *seed* which are easily explained by its limited number of sequences). For RNA families, Infernal seems to accomplish the best predictions in terms of first FP and TPR but nonetheless all the other methods, included HMMer, show comparable results. In fact, although HMMer has the tendency to assign consecutive gaps in the first and last sites of the aligned sequences, these regions are not considered in the comparison, and the core part of the alignment suffices to obtain similar results, in terms of contact prediction, to the other methods. Although in the other metrics presented above there was no clear difference between models learned from large or small seeds, in the contact maps comparison this seems to be an important issue. Indeed, the amount of sequences in the seed slightly affects the quality of the contact map for our methods: for the PF00035 family (whose seed contains only 81 sequences) DCAlign reaches slightly worse performances than HMMer. On the contrary, for the PF00763 all methods produce indistinguishable PPV curves and contact maps. Finally, we remark the results for PF00684 in Fig. 11(b) where DCAlign achieves a better contact prediction, as manifested by the PPV lines. This result, not linked to the way of encoding the seed statistics within the model, but shared by both ℋ_*potts*_ and ℋ_*phmm*_, could be caused by a better treatment of the insertions with respect to HMMer.

**FIG. 11.**
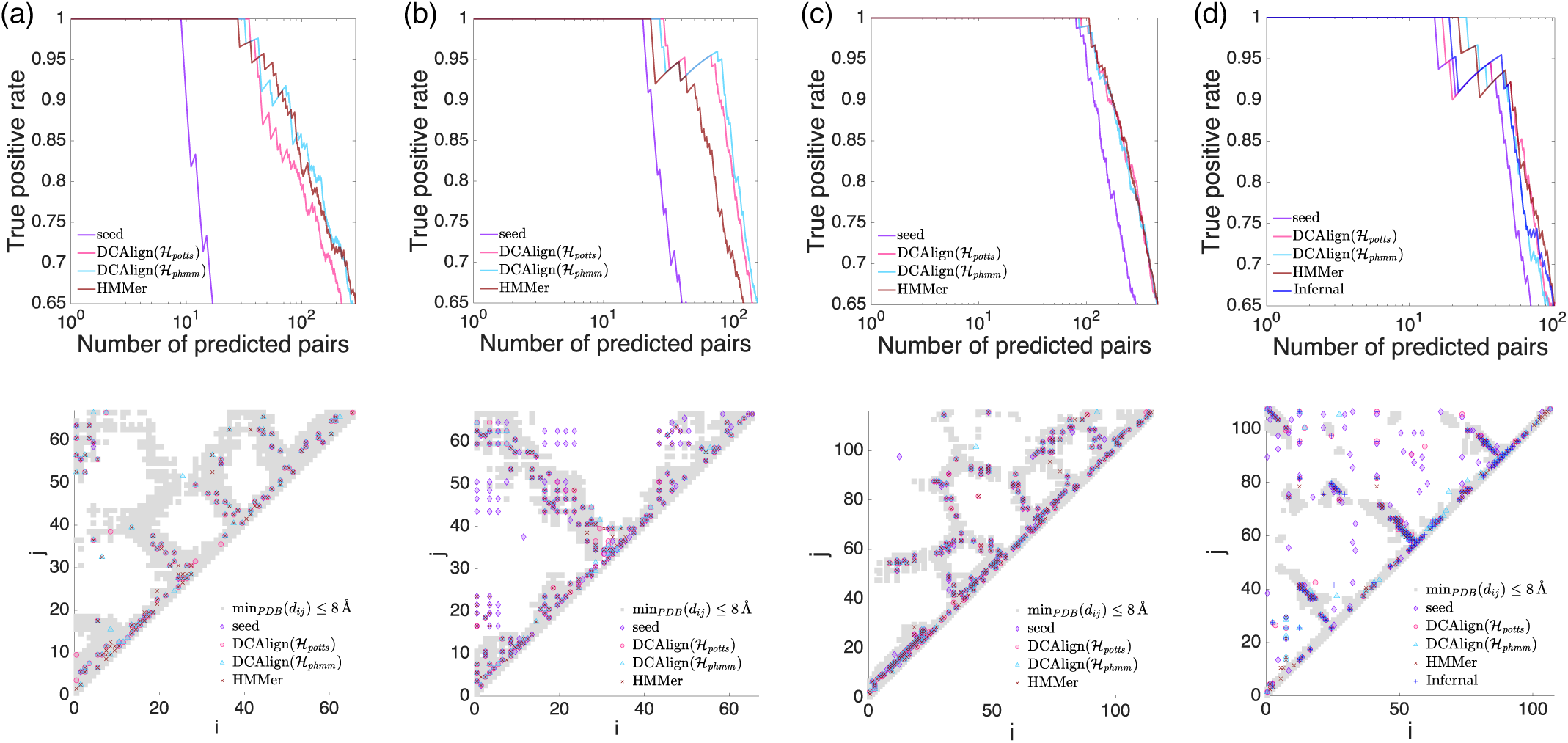
Contact predictions. We show the Positive Predictive Value curves on the top panels and, on the bottom ones, the contact map retrieved by a set of known crystal structures (gray squares) and the Frobenius norms (computed from the full set of aligned sequences or the seed), for (a) PF00035, (b) PF00684, (c) PF00763 and (d) RF00162.

## VIII. CONCLUSIONS

In this work, we have developed and tested DCAlign, a method which allows us to align biological sequences to Potts models of a seed alignment. The set of hyper-parameters characterizing the models are inferred by an inverse statistical-physics based method known as Direct Coupling Analysis from a seed to capture both the single- and two-site (or column) statistics. Single site statistics often signals residue conservation, i.e. the propensity of some sites to restrict the variation of residue (amino-acid or nucleotide) composition because specific residues are functionally and/or structurally important at certain positions. The two-site statistics is related to residue co-variation, or equivalently coevolution. Indeed, residues in direct contact in a folded protein must preserve bio-physically compatible properties, leading to a correlated evolution of pairs of sites.

Most standard alignment algorithms are based on the assumption of independent-site evolution, which is statistically encoded via the so-called profile models. In these alignment procedures, strongly conserved residues serve as anchoring points, and a mismatch in these positions surely induces bad alignment scores (i.e. high energies using a physics-like terminology). Variable sites, characterized by high entropy values in the seed MSA, do provide little information for aligning a new sequence to the seed.

However, often residue pairs show a strong degree of covariation as reflected in two-site statistics and as a consequence, this important collective information must be taken into account. Up to now, the only example in which this information is exploited in the alignment procedure is the case of RNA sequences where the possible base pairing (Watson-Crick or wobble pairs) is encoded in the covariance models used to align.

DCAlign takes advantage of both conservation and co-variation information contained in the seed alignment. It is able to detect within a candidate sequence the most compatible domain among all the possible sub-sequences, as that maximizing a score. The latter can be understood as a probability measure of the domain according to a Boltzmann distribution carefully built from the DCA model and gap/insertion penalties learned from the seed. Using synthetic data at first, we were able to show that the algorithm is able to work under both extreme conditions, when all information is contained in conservation but none in covariation, or vice versa. This renders DCAlign universally applicable, in contrast to more specialized alignment algorithms like HMMer (using profile HMM) or Infernal (using covariance models based on secondary RNA structure).

This universal applicability is well confirmed in the case of real data; we tested both protein and RNA sequence data, aligning large numbers of sequences to the seed MSA provided by the Pfam and Rfam databases. We find that in most cases, our algorithm performs comparably well to the specialized methods, while e.g. profile HMM applied to RNA perform less well. Interestingly, in one of the studied protein families, we find a large group of proteins, which are aligned differently by HMMer and DCAlign. The sequences aligned by DCAlign show a better coherence with the sequence-space structure of the seed MSA than those aligned by HMMer, suggesting that the alignment proposed by DCAlign is actually to be preferred in this case.

## ACKNOWLEDGMENTS

The authors thank Sean Eddy, Alessandra Carbone, Francois Coste and Hugo Talibart for interesting discussions. APM thanks Edoardo Sarti for interesting discussions and his assistance with the *Pfam interactions* code. APM, AP, and MW acknowledge funding by the EU H2020 research and innovation programme MSCA-RISE-2016 under grant agreement No. 734439 INFERNET. APM and FZ acknowledge funding from the Simons Foundation (#454955, Francesco Zamponi) and the access to the HPC resources of MesoPSL financed by the Region Ile de France and the project Equip@Meso (reference ANR-10-EQPX-29-01) of the programme Investissements d’Avenir supervised by the Agence Nationale pour la Recherche.

## SUPPORTING INFORMATION

### 1. Belief propagation equation

In order to write the BP equations, it is convenient to introduce weights *𝒲*_*i,j*_ associated to pair interactions in the Boltzmann weight Eq. (15). For *i < j* we define

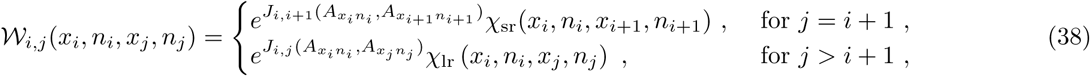

and for *i > j* we just set *𝒲*_*i,j*_(*x*_*i*_, *n*_*i*_, *x*_*j*_, *n*_*j*_) = *𝒲*_*j,i*_(*x*_*j*_, *n*_*j*_, *x*_*i*_, *n*_*i*_). The total Boltzmann weight then becomes

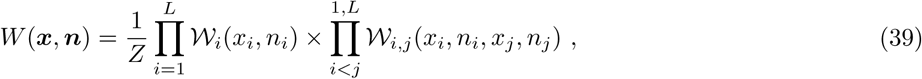

and the BP iteration equations are then written straightforwardly:

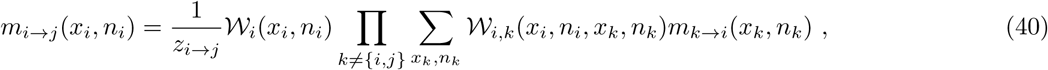

where *m*_*i*→*j*_(*x*_*i*_, *n*_*i*_) are the BP messages. From the converged messages, the marginal probability of node *i* is estimated as

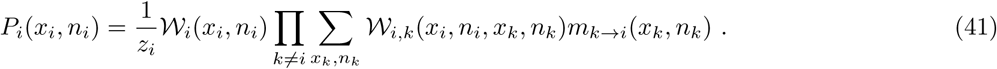

Note that if all the couplings vanish for |*i* − *j*| *>* 1 and the *χ*_lr_ are omitted, then *𝒲*_*ij*_ = 1 for |*i* − *j*| *>* 1 and the BP equations reduce to the transfer matrix equations, which are exact in that case. In presence of long range couplings, instead, we stress that the *χ*_lr_ are redundant and can be omitted in an exact treatment. However, the approximate BP equations depend on the *χ*_lr_ and give different results if these terms are omitted. In other words, the BP approximation does not “commute” with the insertion/removal of the *χ*_lr_ constraints.

### 2. From belief propagation to mean field

We can obtain the mean field equations by considering a weak interaction (or large connectivity) limit of the BP equations. This limit consists in making two approximations for pairs of sites *j* ≠ {*i* + 1, *i* − 1} that are not nearest-neighbor in the linear chain:

1. We assume that *J*_*i,j*_(*A, B*) is sufficiently small to approximate 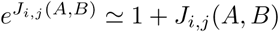. This is likely the case for far away sites that are not in contact in the three-dimensional protein structure. Moreover, when the Hamiltonian is written in zero-sum gauge, all couplings tend to be small (often the largest ones are ∼1), because this gauge choice minimizes the Frobenius norm of the coupling matrices *J*_*i,j*_.
2. We approximate the messages *m*_*i*→*j*_(*x*_*i*_, *n*_*i*_) with the marginal density *P*_*i*_(*x*_*i*_, *n*_*i*_), which is a reasonable choice when *i* and *j* and far enough that the influence of *j* on *i* is negligible.

We emphasize that while both approximations are exact in a mean field setting, in which one takes the thermodynamic limit *L* with couplings vanishing with *L*→ *∞*, they do not hold exactly for our setting in which both *L* and the couplings are finite. Moreover, an important remark is that this approximation does not preserve the gauge invariance of the Hamiltonian.

To see the effect of these approximations, consider, for instance, the marginal density of a node 1 *< i < L* in the belief propagation framework. Let us call the nearest-neighbor messages *F*_*i*_(*x*_*i*_, *n*_*i*_) = *m*_*i*→*i*+1_(*x*_*i*_, *n*_*i*_) and *B*_*i*_(*x*_*i*_, *n*_*i*_) = *m*_*i*→*i*−1_(*x*_*i*_, *n*_*i*_). The contributions coming from the right and from the left along the linear chains are then

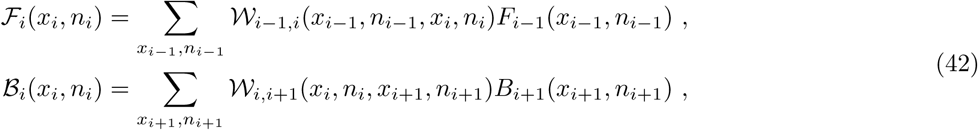

and we have

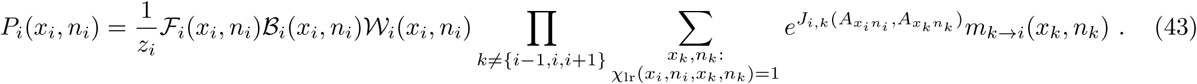

Applying the weak interaction approximation to the contribution of distant sites, and introducing

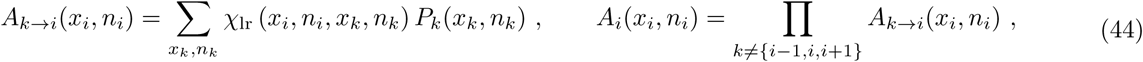

we obtain

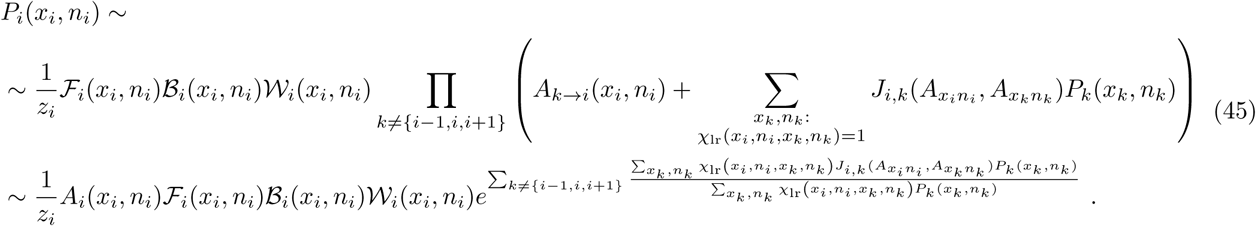

Introducing the modified weight

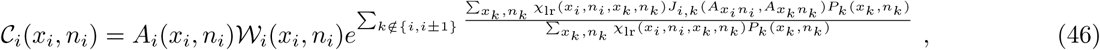

we obtain that

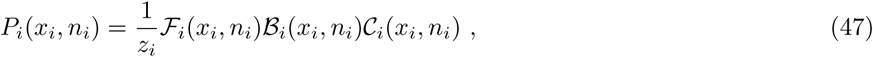

which amounts to use the transfer matrix expression with the replacement *𝒲*_*i*_ → *𝒞*_*i*_. A very similar procedure can be applied to treat the long range part in the forward and backward messages, *F*_*i*_ and *B*_*i*_, with the same result: the equations are identical to the transfer matrix equations with *𝒲*_*i*_ →*𝒞*_*i*_. This procedure thus provides a set of closed equations for the forward and backward messages and the marginal probabilities.

For simplicity, we also make an additional approximation, namely that for each pair of distant sites *i, k* and for each “relevant” choice (in a sense that will be more precise below) of *x*_*i*_, *n*_*i*_, we have

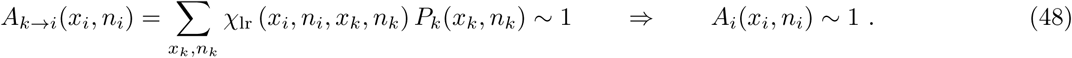

In words, this amounts to assume that the long-range constraints we artificially introduce do not play a big role on the normalization of marginals, i.e. that most of the mass of *P*_*k*_ is concentrated on compatible assignments of *x*_*k*_, *n*_*k*_. Note that for very unlikely values of *x*_*i*_, *n*_*i*_ (e.g. assigning *x*_*i*_ = 0, *n*_*i*_ = *N* + 1 at the very beginning of the sequence), the approximation in Eq. (48) is probably not correct. However, the probability *P*_*i*_ for such values is already suppressed by the short-range terms, making the error irrelevant. Under this approximation, the modified weights reduce to

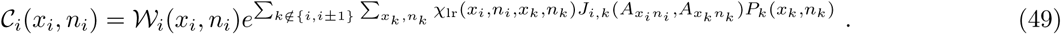

The advantage of this additional approximation is that when the long-range couplings *J*_*i,j*_ → 0, the mean field equations reduce exactly to the transfer matrix equations, as it should be.

### 3. Mean field free energy

We give here the expression of the free energy associated with the Boltzmann weight in Eq. (15). This free energy can be used, for example, as a score to compare the quality of the alignment of different sub-regions of the same very long sequence. The free energy associated with Belief Propagation is given by

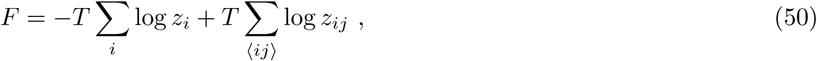

where the second sum is over distinct pairs ⟨*ij*⟩. Here, the site term *z*_*i*_ is the denominator in Eq. (41), while the link term is given by

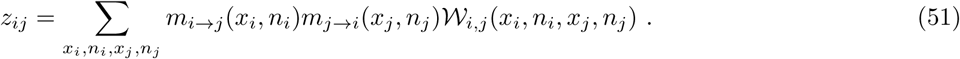

In the weak interaction approximation, we can write the site term *z*_*i*_ as simply the denominator of the single-site marginals, as defined in Eq. (47), i.e.

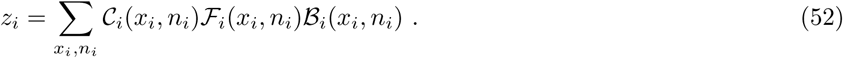

(with small differences on the boundaries). For the link term *z*_*ij*_ we should distinguish between nearest neighboring sites, and far away sites. For the first case we get

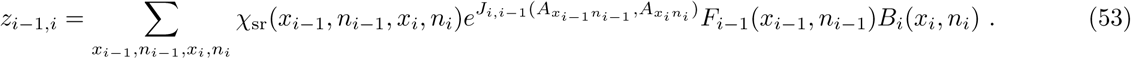

For the second case, *j ≠ i ±* 1, we get,

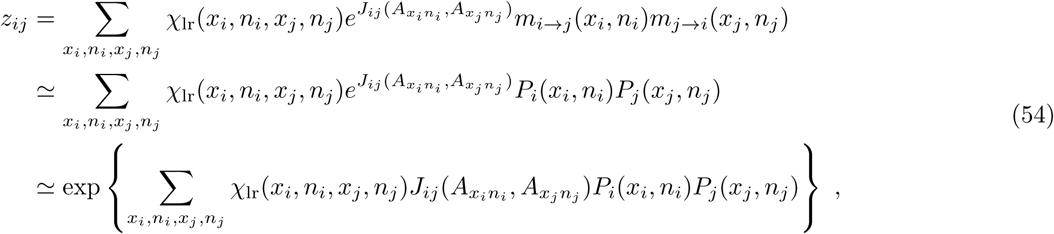

where we applied the mean field approximation of identifying messages with marginals (first to second line), and considering the interactions *J*_*ij*_ as being small (second to third line). Note that in the second step we assumed that 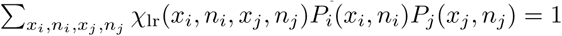, i.e. that the marginals respect the long-range constraints.

### 4. Zero temperature limit

We discuss here briefly how to take the zero temperature limit of the mean field equations. In order to introduce a temperature *T* = 1*/β ≠* 1 we need to rescale all the parameters in the cost function, as *J*_*ij*_ → *βJ*_*ij*_, *h*_*i*_ → *βh*_*i*_, ***µ*** → *β****µ, λ*** → *β****λ***.

In order to take the *T* → 0 limit we define

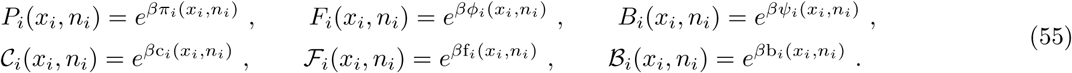

We also define 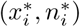 the maximum of *π*(*x*_*i*_, *n*_*i*_). Note that in the first line the messages are normalized, so 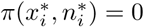, and a similar relation for the other messages. In the second line instead, messages are not normalized. The zero temperature mean field equations are then obtained by taking the limit *β* → *∞*, in which the sums over *x*_*i*_, *n*_*i*_ are dominated by the maximum of the integrand. One should only take into account that the hard constraints *χ*_in_ and *χ*_end_ set to zero some elements of *P*_1_, *F*_1_, *P*_*L*_ and *B*_*L*_, which translates in −*∞* elements for *π*_1_, *ϕ*_1_, *p*_*L*_ and *?*_*L*_. We obtain (with minor modifications at the boundaries):

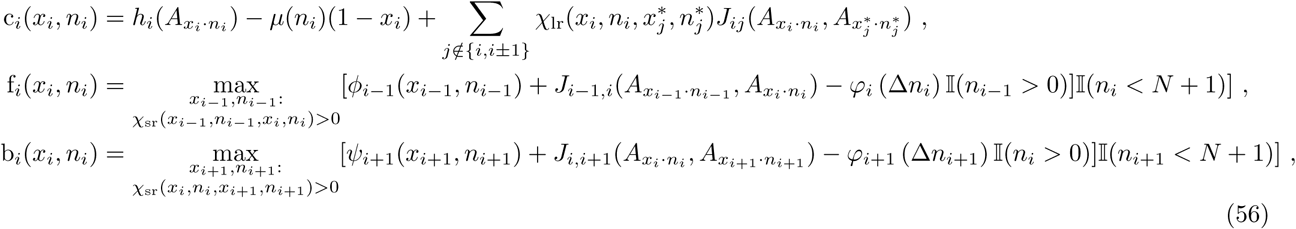

together with

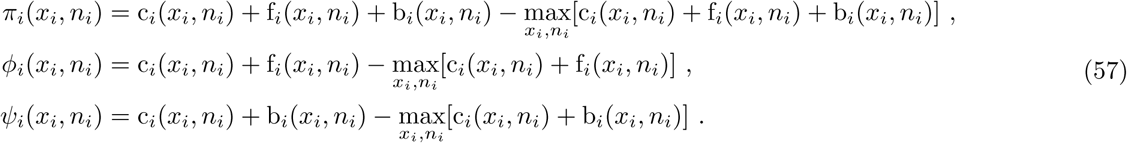

### 5. Maximum likelihood equations for insertions

We determine the values of the insertion penalties *λ*_*o*_, *λ*_*e*_ by maximizing the likelihood of the data, i.e. the *M* sequences of the seed, given the parameters:

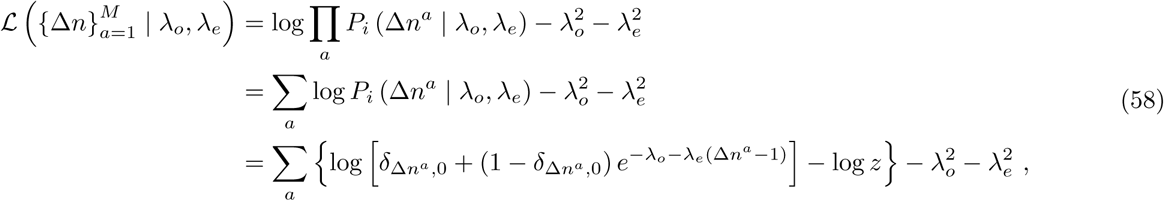

where the regularization terms are used to avoid infinite of undetermined parameters. From the likelihood we obtain an explicit expression of the gradient,

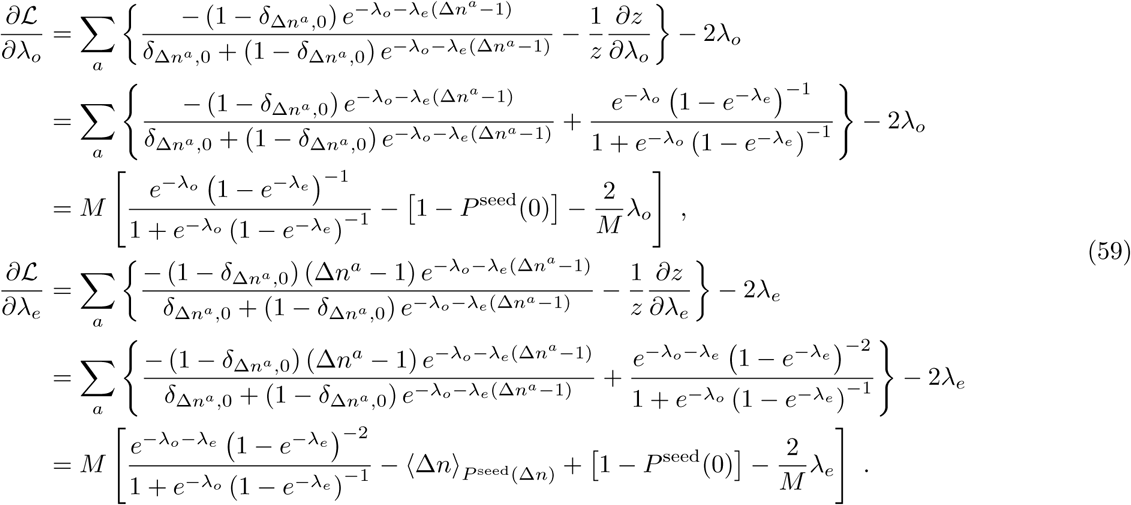

The maximum likelihood estimators can then be obtained by gradient descent, as detailed in the main text.

### 6. Sequence-based distances plot

**FIG. 12.**
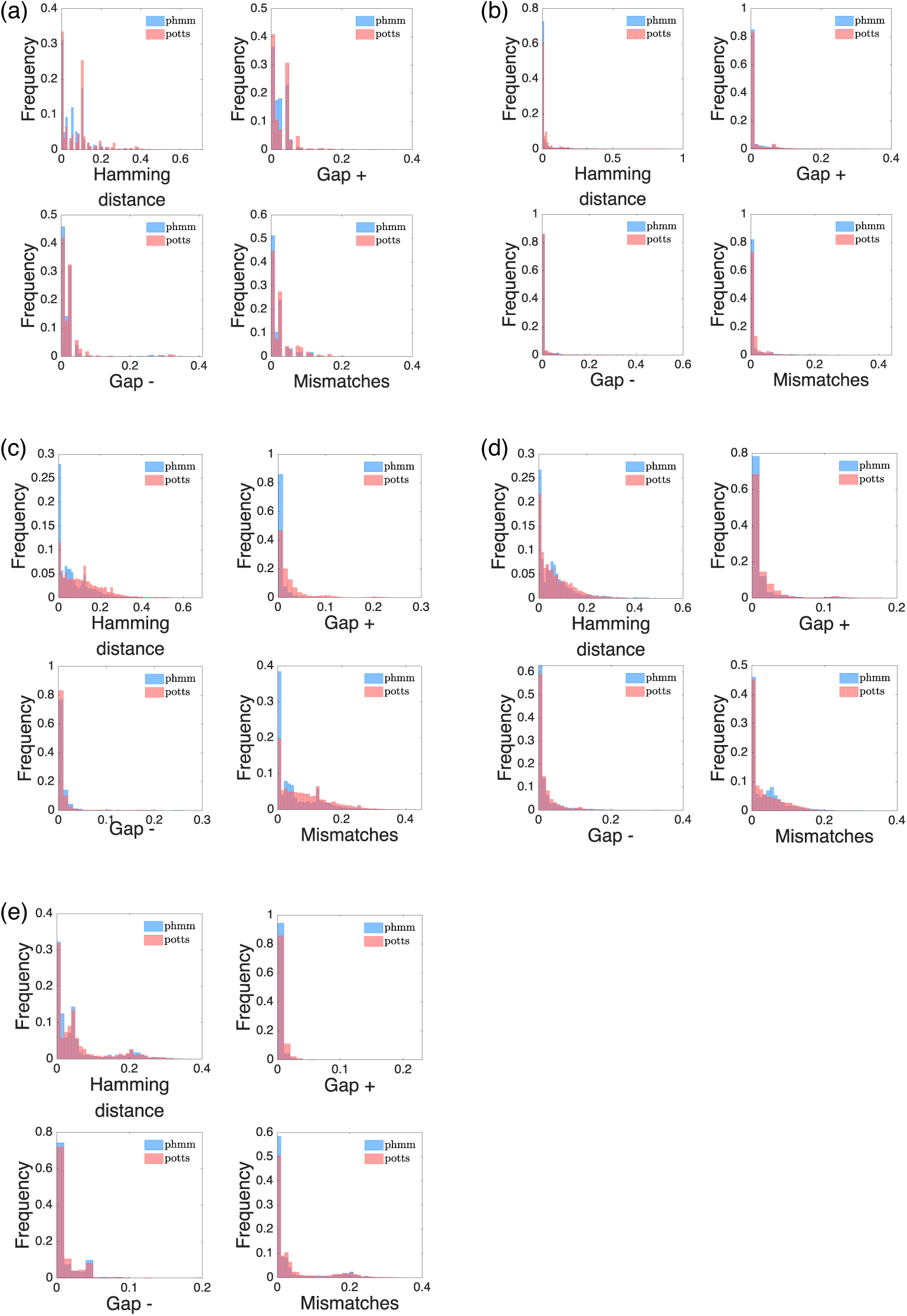
Distances distribution. We plot here the histograms of the Hamming distances, Gap_+_, Gap_−_ and Mismatches for the protein families PF00684 (a), PF00763 (b) using as reference the HMMer results and as target the DCAlign results, and for RF00059 (c), RF00167 (d) and RF01734 (e) using as reference the Infernal results and as target the DCAlign results.

### 7. Proximity measures plot

**FIG. 13.**
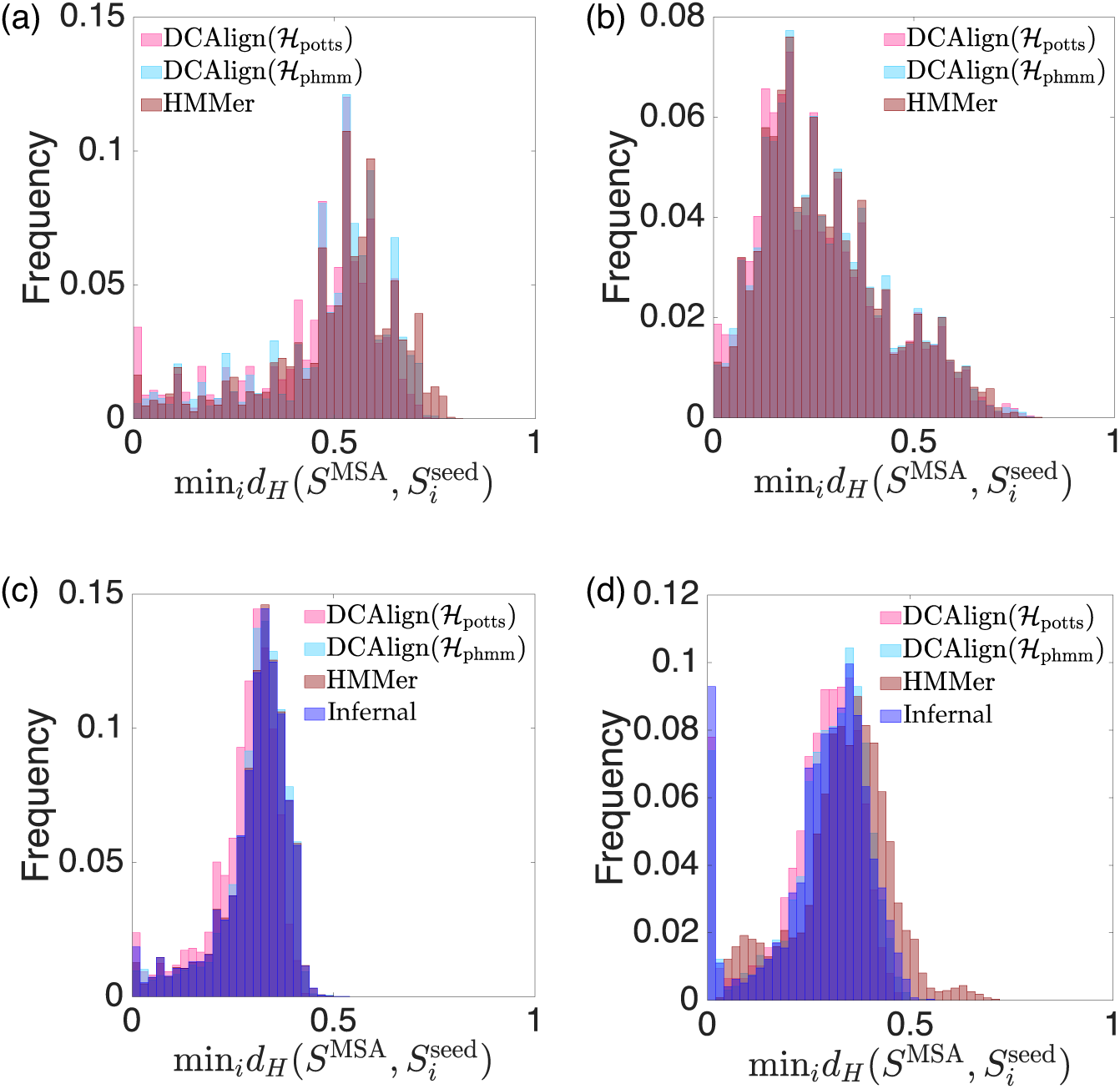
Proximity measures. Histograms of the minimum distances computed according to Eq. (32) for the full set of aligned sequences obtained by DCAlign-*potts*, DCAlign-*phmm*, HMMer, and Infernal, against the seed. Panels (a), (b), (c) and (d) refer to the families PF00035, PF00763, RF00059, RF00167 respectively.

### 8 Contact maps

**FIG. 14.**
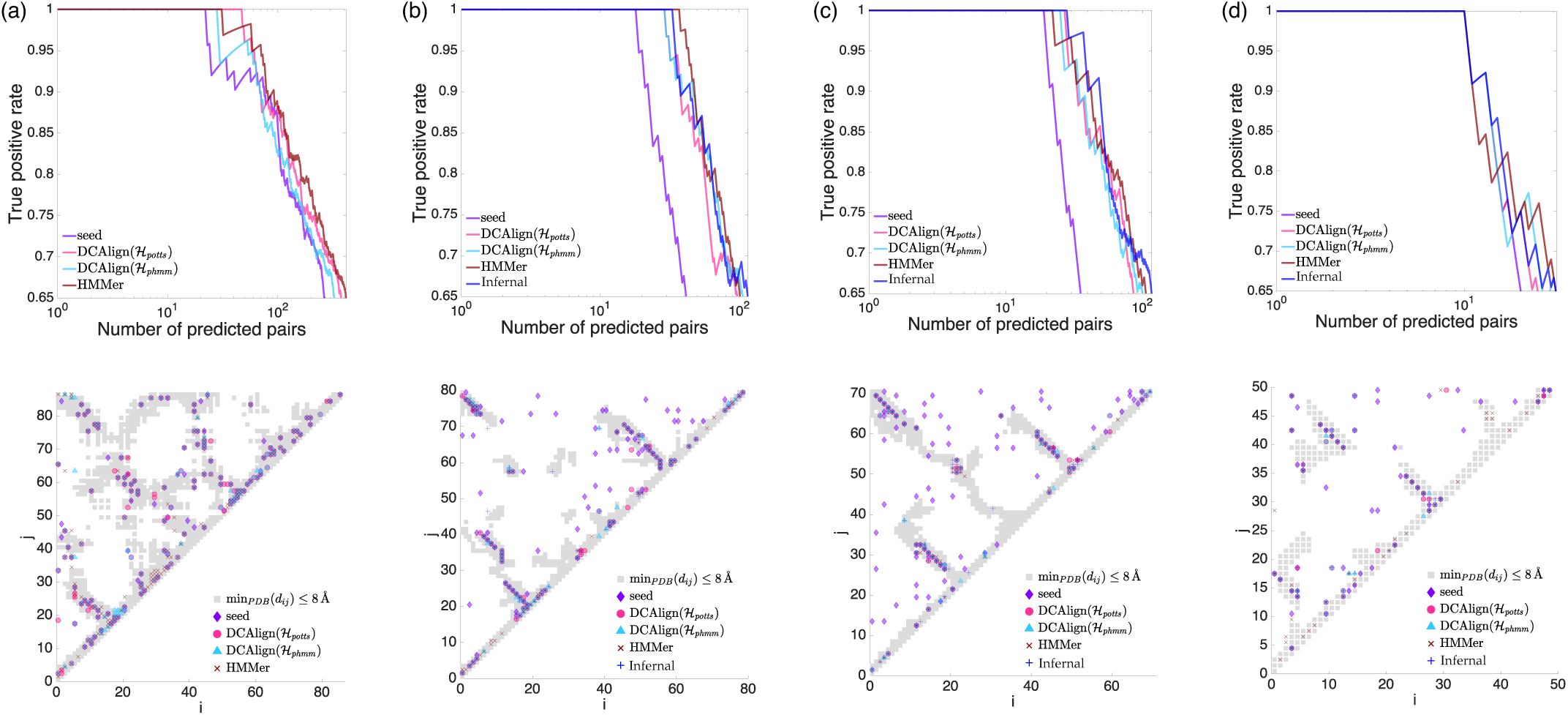
Contact predictions. We show the Positive Predictive Value curves on the top panels and, on the bottom ones, the contact map retrieved by a set of known crystal structures (gray squares) and the Frobenius norms (computed from the full set of aligned sequences or the seed), for (a) PF00677, (b) RF00059, (c) RF00167 and (d) RF01734.

